# Phosphoregulated SMCR8-FIP200 interaction connects the ALS/FTD-linked C9orf72 complex to autophagy initiation and mitochondrial quality control

**DOI:** 10.64898/2026.04.16.718395

**Authors:** Jian Wang, Colin Davis, Simone Kunzelmann, Sarah Maslen, Gavin Kelly, Mark Skehel, Anne Schreiber

**Affiliations:** Cellular Degradation Systems Laboratory, The Francis Crick Institute, 1 Midland Road, NW1 1AT London, UK; Structural Biology STP, The Francis Crick Institute, 1 Midland Road, NW1 1AT London, UK; Proteomics STP, The Francis Crick Institute, 1 Midland Road, NW1 1AT London, UK; Bioinformatics and Biostatistics STP, 1 Midland Road, NW1 1AT London, UK

## Abstract

Hexanucleotide (GGGGCC) repeat expansions in the non-coding region of *C9ORF72* are a major genetic cause of ALS/FTD and reduce C9orf72-SMCR8-WDR41 complex levels, but how this contributes to autophagy-lysosome dysfunction and previously reported mitochondrial quality-control defects in C9ORF72-ALS/FTD remains unclear. Here we identify a direct interaction between SMCR8 and the FIP200 subunit of the ULK1/2 autophagy initiation complex, mediated by two FIP200-interacting region (FIR) motifs in a disordered SMCR8 loop. Phosphorylation of these motifs by ULK1/2 or TBK1 strengthens binding and promotes ULK1/2 complex association in cells. Stabilising the SMCR8-FIP200 interaction suppresses Parkin-dependent mitophagy, whereas both stabilisation and weakening impair deferiprone-induced mitophagy, while leaving bulk autophagy, lysophagy and ivermectin-induced mitophagy largely intact. These findings define a regulated C9orf72–ULK1/2 axis and provide a mechanistic framework by which repeat-expansion-associated reduction in C9orf72 complex abundance may contribute to previously observed mitochondrial quality-control defects in C9ORF72-ALS/FTD.

## Introduction

Amyotrophic lateral sclerosis (ALS) is a rapidly progressive motor neuron disease characterised by loss of voluntary muscle control and, ultimately, fatal respiratory failure. Frontotemporal dementia (FTD) primarily affects the frontal and temporal lobes, leading to profound behavioural changes and cognitive decline. Despite their distinct clinical presentation, ALS and FTD lie on a pathological and genetic continuum. A GGGGCC repeat expansion (G_4_C_2_)_n_ in a non-coding region of the *C9ORF72* gene is the most frequent genetic cause of familial ALS and FTD, accounting for up to ∼40% of cases^1,2^. The expansion drives disease through multiple mechanisms, including toxic gain-of-function effects (repeat RNA foci with sequestration of RNA-binding proteins^3–7^, and dipeptide repeat proteins produced by repeat-associated non-AUG translation^8–12)^. In addition, reduced *C9ORF72* mRNA or C9orf72 protein levels have been observed in patient-derived cells and postmortem tissue^13–15^, suggesting that reduced C9orf72 protein and as a result C9orf72 complex levels may contribute to disease. Defining the physiological function of the C9orf72 complex is therefore central to understanding how reduced C9orf72 levels might intersect with cellular vulnerability in ALS/FTD.

At the molecular level, C9orf72 forms a heterotrimeric complex with SMCR8 and WDR41 (hereafter “the C9orf72 complex” or “CSW”)^16–19^. Both C9orf72 and SMCR8 contain DENN-family domains, which are frequently found in regulators of small GTPases, and the C9orf72 complex has been linked to Rab- and Arf-family pathways that control membrane trafficking and organelle organisation^17,19–28^. In line with these functions, the C9orf72 complex also participates in lysosomal nutrient sensing: upon amino acid depletion, WDR41 promotes recruitment of the complex to lysosomes through interactions with lysosomal components, including the transporter PQLC2, thereby contributing to restoration of mTORC1 signalling upon nutrient refeeding^29,30^.

Macroautophagy (hereafter “autophagy”) is a conserved intracellular degradation pathway that maintains cellular homeostasis and promotes quality control, cell-intrinsic defence and stress adaptation by delivering cytosolic material to lysosomes for degradation and recycling. During nutrient starvation, autophagy can proceed as a non-selective process (bulk autophagy) in which portions of the cytoplasm are sequestered within double-membrane vesicles (autophagosomes) for lysosomal turnover^31^. Autophagy also comprises selective branches that target defined cargos, including protein aggregates, intracellular pathogens and damaged organelles, such as lysosomes (lysophagy) and mitochondria (mitophagy)^32^. Selectivity is mediated by cargo adaptors that recognise autophagy cargos, often via ubiquitin or organelle-specific signals, and recruit the core autophagy machinery to promote local autophagosome biogenesis^33–37^. Notably, autophagy-lysosome pathway dysfunction is widely reported in ALS/FTD and is genetically linked to disease through mutations in genes encoding key selective autophagy regulators, including TANK-binding kinase 1 (TBK1) and the cargo adaptors Optineurin (OPTN) and p62^38,39^, underscoring the importance of defining how ALS/FTD-associated pathways intersect with the autophagy machinery.

In mammalian cells, autophagy initiation is orchestrated by the ULK1/2 complex, composed of the serine/threonine kinases ULK1 or ULK2, the scaffold subunit FIP200, ATG13 and ATG101^40–44^. Upon clustering and activation, ULK1/2 drives autophagosome biogenesis by coordinating recruitment of ATG9A-positive vesicles and activating the class III phosphatidylinositol 3-kinase complex I (PI3KC3-C1) to generate PI3P^31,45–50^. PI3P then recruits WIPI proteins^51^ which in turn recruit the ATG12–ATG5-ATG16L1 complex^52^ to mediate LC3/GABARAP-family protein lipidation to promote autophagosome formation and subsequent fusion with lysosomes^53–56^.

In selective autophagy, FIP200 plays a central role as a recruitment hub. Many cargo adaptors engage the C-terminal Claw domain of FIP200^57–60^, whereas others bind a second site within the coiled-coil region^61,62^. Claw domain recognition is mediated by short linear FIP200-interacting region (FIR) motifs present in many cargo adaptors, as well as in autophagy regulators and core autophagy proteins^63,64^, thereby coupling cargo recognition to local ULK1/2 recruitment and autophagosome biogenesis.

A growing body of work has linked the C9orf72 complex to autophagy and lysosome biology, yet key mechanistic questions remain unresolved. Several studies support a model in which the C9orf72 complex promotes autophagy and lysosomal homeostasis, with C9orf72 depletion impairing autophagy in multiple systems, including neuronal models^65–67^. Loss of SMCR8 has likewise been reported to impair autophagy^19,68^. However, the literature is not fully consistent, as increased autophagic flux after *C9orf72* knockdown has also been described^19,69^, highlighting the complexity of interpreting knockout or knockdown phenotypes for a protein complex that additionally functions in lysosomal amino acid sensing, mTORC1-linked lysosome homeostasis and membrane trafficking^70–73^. Moreover, the C9orf72 complex has been reported to associate with the ULK1/2 autophagy initiation machinery and with selective autophagy regulators, including TBK1 and the cargo adaptor p62^17–20,74–76^, but which of these connections are direct, how they are regulated, and which autophagy branches they control has remained unclear.

Here, we define a direct, phosphorylation-regulated molecular link between the C9orf72 complex and the autophagy initiation machinery. Using purified recombinant proteins, we show that the C9orf72 complex directly binds FIP200 and that this interaction is strongly enhanced upon phosphorylation by ULK1/2 and TBK1. We map binding to the C-terminal Claw domain of FIP200 and show that the SMCR8 subunit mediates this interaction via two phosphoregulated FIR motifs embedded within a long intrinsically disordered loop. Quantitative binding measurements support an avidity-driven mechanism in which these two motifs enable high-affinity binding to the dimeric FIP200 Claw domain. In cells, SMCR8 separation-of-function mutants that selectively weaken or stabilise FIP200 binding tune ULK1/2 complex association and result in stimulus-dependent effects on selective autophagy that are most evident across distinct mitophagy pathways, while leaving bulk autophagy and lysophagy largely intact. These findings indicate that regulated SMCR8-dependent FIP200 engagement is particularly important for specific mitochondrial quality control pathways. Together, these findings define a regulated C9orf72-ULK1/2 complex axis linking SMCR8-dependent FIP200 engagement to mitophagy and provide a plausible mechanistic basis by which repeat-expansion-associated reduction in C9orf72 complex abundance could contribute to previously observed mitochondrial quality-control defects in C9ORF72-ALS/FTD^77–79^.

## Results

### The C9orf72 complex directly interacts with FIP200 and ULK1 in a phosphorylation-dependent manner

The C9orf72 complex has previously been reported to interact with the ULK1/2 complex, as well as with proteins involved in selective autophagy, including the TBK1 kinase and the cargo adaptor Optineurin (OPTN)^17–20,74,75^. To test whether these interactions are direct and occur in the context of the intact C9orf72 complex (abbreviated as “CSW” – <u>C</u>9orf72/<u>S</u>MCR8/<u>W</u>DR41 – in all figures to clearly distinguish the complex from the C9orf72 subunit), we expressed and purified recombinant C9orf72 complex (**Fig. 1a** and **Extended Data Fig. 1a**) together with candidate autophagy-related interactors for *in vitro* pulldown assays (**Fig. 1b**). For the ULK1/2 complex, we included the individual subunits ULK1 (both full length and kinase domain only), ULK2, and FIP200, together with the ATG13-ATG101 subcomplex. We also tested ULK3, an ULK family homologue whose role in canonical ULK1/2-driven autophagy initiation is less well defined, as well as the cargo adaptors OPTN and TAX1BP1 and the LC3/GABARAP-specific E3 ligase ATG12–ATG5-ATG16L1, which has previously been reported to associate with the ULK1/2 complex^64,80^ and could therefore mediate interactions with the C9orf72 complex. In addition, we also included the TBK1 kinase in complex with its autophagy-related adaptor 5-azacytidine-induced protein 2 (AZI2; also known as NAP1)^81^. Strikingly, the recombinant C9orf72 complex directly interacted with full-length FIP200 and also displayed weak affinity for ULK1, but not for the isolated ULK1 kinase domain (ULK1^1–283^) or the close homologues ULK2 and ULK3 (**Fig. 1c, d**). Under the tested conditions, the C9orf72 complex also showed no appreciable binding to TBK1-NAP1, ATG13-ATG101, the ATG12–ATG5-ATG16L1 complex, or the cargo adaptors OPTN and TAX1BP1 (**Fig. 1c**).

**Fig. 1.**
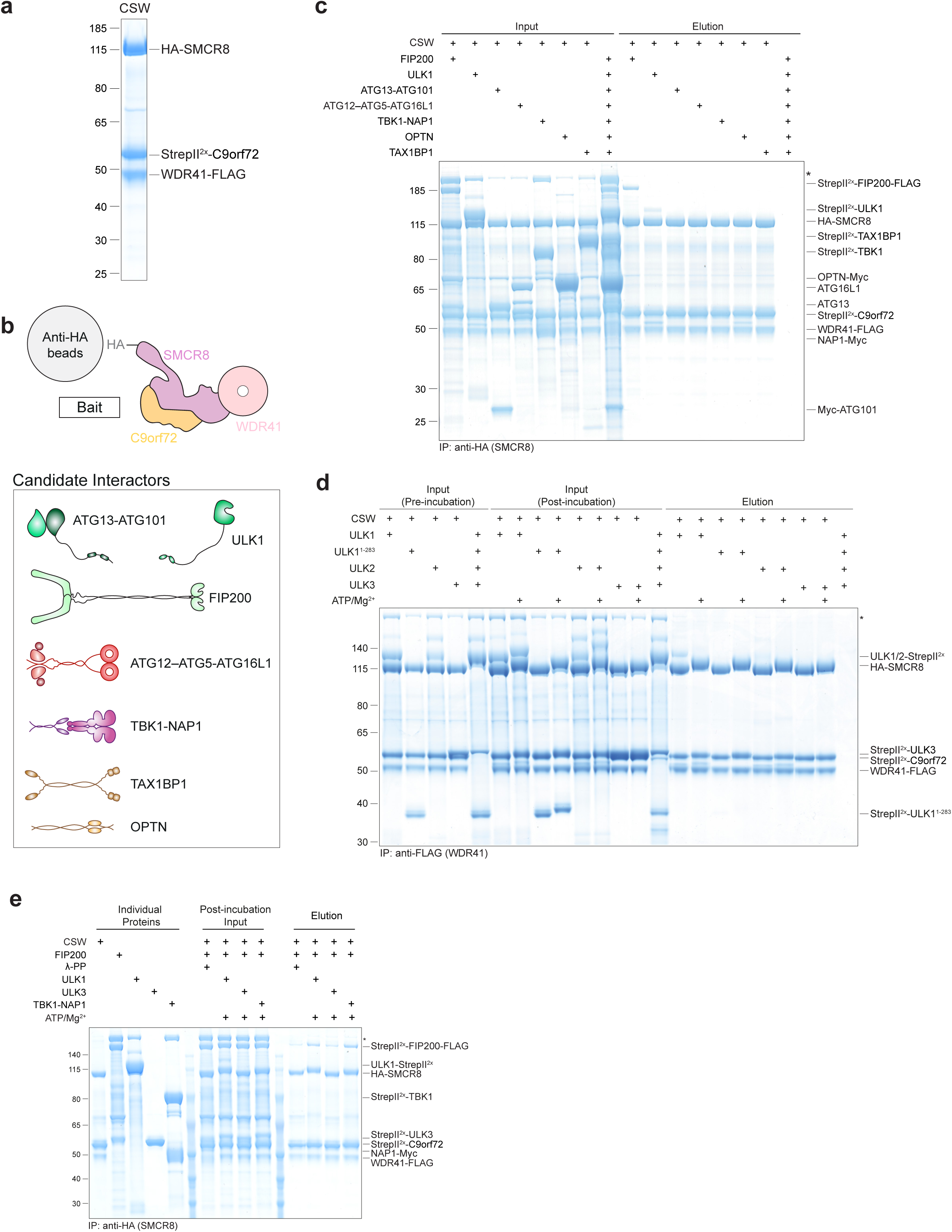
The C9orf72 complex binds directly to FIP200 and ULK1 in a phosphorylation dependent manner. **a**, Purification of recombinant *H. sapiens* C9orf72 complex (CSW). The C9orf72 complex was purified by StrepTactin affinity chromatography followed by size-exclusion chromatography. A representative SEC peak fraction is shown. Molecular weight markers (kDa) are indicated on the left. **b**, The C9orf72 complex, containing HA-tagged SMCR8, was used as bait to test for direct interactions with FIP200, ULK1, ATG13-ATG101, ATG12–ATG5-ATG16, TBK1-NAP1, OPTN and TAX1BP1.**c**, The C9orf72 complex directly interacts with FIP200 and ULK1. The recombinant C9orf72 complex containing HA-tagged SMCR8 was immobilised on anti-HA agarose and incubated with the proteins shown in Fig. 1b. After washing, bound proteins were eluted, and input and elution fractions were analysed by SDS-PAGE followed by Coomassie staining. An asterisk (*) denotes an endogenous insect-cell protein that binds to Strep-Tactin resin and co-elutes with Strep-tagged proteins. **d**, The interaction between the C9orf72 complex and ULK1 is negatively regulated by ULK1-mediated phosphorylation. The recombinant C9orf72 complex was immobilised on anti-FLAG agarose and incubated with full-length ULK1, the ULK1 kinase domain (ULK1^1–283^), ULK2 or ULK3 in the presence or absence of ATP/Mg²⁺. After washing, bound proteins were eluted, and input (pre- and post-incubation) and elution fractions were analysed by SDS-PAGE followed by Coomassie staining. **e**, The interaction between FIP200 and the C9orf72 complex is strengthened by ULK1- and TBK1-mediated phosphorylation. FIP200 and the recombinant C9orf72 complex were combined and either dephosphorylated with lambda protein phosphatase (λ-PP) or phosphorylated with recombinant ULK1, ULK3 or TBK1 (in complex with NAP1) at a highly substoichiometric ratio (1:20 relative to the C9orf72 complex) in the presence of protein phosphatase inhibitors. The C9orf72 complex was captured on anti-HA agarose, and after washing, bound proteins were eluted. Input and elution fractions were analysed by SDS-PAGE followed by Coomassie staining.

The ULK1 kinase is a central regulator of autophagy that phosphorylates numerous autophagy-related proteins to drive autophagosome formation^46,82–88^. We therefore tested whether ULK1-mediated phosphorylation modulates C9orf72 complex binding to ULK1 or FIP200. Strikingly, while ULK1-mediated phosphorylation abolished binding of ULK1 to the C9orf72 complex (**Fig. 1d**), it significantly enhanced the interaction between the C9orf72 complex and FIP200 (**Fig. 1e**). As the C9orf72 complex has been reported to interact with the TBK1 kinase^18–20,74,75^, we next tested whether TBK1 can also regulate the interaction between FIP200 and the C9orf72 complex. Indeed, when we compared the effects of ULK1-, TBK1-, and ULK3-dependent phosphorylation, both ULK1 and TBK1 enhanced the FIP200-C9orf72 complex interaction, whereas ULK3 had no noticeable effect (**Fig. 1e**).

It has been shown that the C9orf72 complex-ULK1/2 complex interaction is enhanced upon bulk autophagy induction^19^, a condition under which ULK1 is activated^46^. Because ULK1-mediated phosphorylation selectively strengthens the interaction between the C9orf72 complex and FIP200 (**Fig. 1e**), but not its interaction with ULK1 itself (**Fig. 1d**), we focused subsequent analyses on the direct C9orf72 complex-FIP200 interaction to define its molecular basis and assess its physiological relevance in autophagy.

### The C9orf72 complex directly binds the FIP200 Claw domain

To map the C9orf72 complex binding site on FIP200, we tested several FIP200 truncation mutants: the N-terminal dimerization domain (FIP200^NTD^), the central region comprising the intrinsically disordered region (IDR) and most of the coiled-coil (CC) domain (FIP200^CR^, amino acids 641-1329), and the C-terminal region (FIP200^CTR^, amino acids 1330-1594) (**Fig. 2a**). Notably, only the FIP200^CTR^ construct, which includes the Claw domain and an adjacent CC segment encompassing the NDP52/TAX1BP1-binding sites^61,89^, showed specific binding to the C9orf72 complex (**Fig. 2b**). Further dissection of the FIP200^CTR^ demonstrated that binding is mediated by the FIP200 Claw domain (FIP200^Claw^, aa 1494–1594), rather than by the preceding CC region (FIP200^1330–1493^) (**Fig. 2c**).

**Fig. 2.**
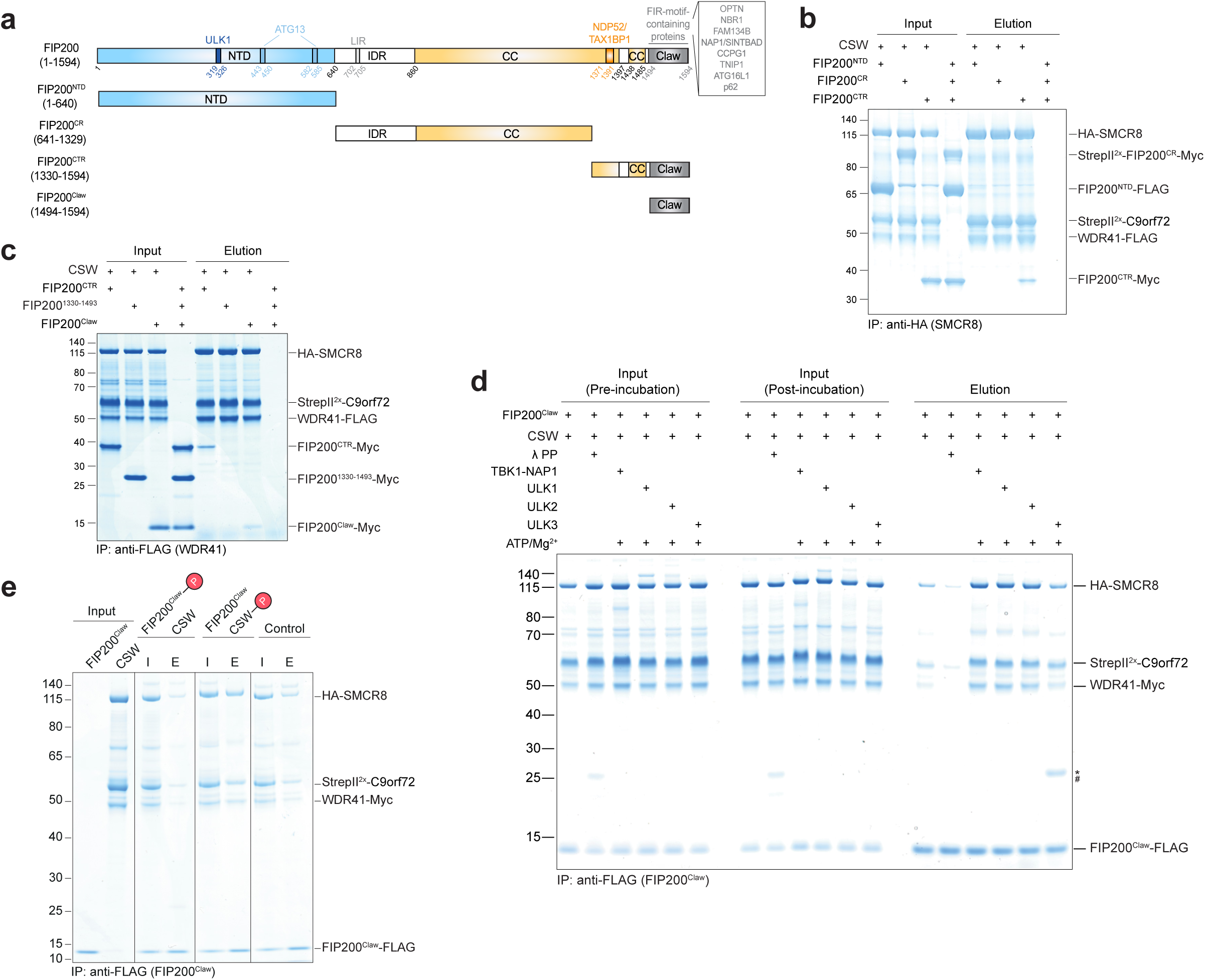
The C9orf72 complex directly binds the FIP200 Claw domain, and kinases central to autophagy enhance this interaction by phosphorylating the C9orf72 complex. **a**, Domain organisation of FIP200 and truncation constructs used for interaction mapping. Schematic of FIP200 showing the N-terminal dimerization domain (FIP200^NTD^; aa 1–640), the central region (FIP200^CR^; aa 641–1329) comprising the intrinsically disordered region (IDR) and most of the coiled-coil (CC) domain, and the C-terminal region (FIP200^CTR^; aa 1330–1594) containing the Claw domain (aa 1494–1594). **b**, The C9orf72 complex interacts directly with the FIP200^CTR^. The recombinant C9orf72 complex (CSW) containing HA-tagged SMCR8 was immobilised on anti-HA agarose and incubated with the FIP200^NTD^, FIP200^CR^ or FIP200^CTR^. After washing, bound proteins were eluted, and input and elution fractions were analysed by SDS-PAGE followed by Coomassie staining. **c**, C9orf72 complex binding is mediated by the FIP200 Claw domain. The recombinant C9orf72 complex containing FLAG-tagged WDR41 was immobilised on anti-FLAG agarose and incubated with the FIP200^CTR^, FIP200^1330–1493^ or the FIP200 Claw domain (FIP200^Claw^). After washing, bound proteins were eluted, and input and elution fractions were analysed by SDS-PAGE followed by Coomassie staining. **d**, The C9orf72 complex directly binds the FIP200 Claw domain, and this interaction is strengthened by ULK1/2- and TBK1-mediated phosphorylation. The FIP200 Claw domain (FIP200^Claw^) and C9orf72 complex (CSW) were combined and left either untreated, dephosphorylated with λ-PP, or phosphorylated with recombinant ULK1, ULK2, ULK3 or TBK1 (in complex with NAP1) at a highly substoichiometric ratio (1:20 relative to the C9orf72 complex) in the presence of protein phosphatase inhibitors. FIP200^Claw^ was captured on anti-FLAG agarose, and after washing, bound proteins were eluted. Input (pre- and post-incubation) and elution fractions were analysed by SDS-PAGE followed by Coomassie staining. Hash (#) and asterisk (*) indicate two closely migrating bands corresponding to λ-PP and the IgG light chain, respectively. **e**, ULK1-mediated phosphorylation of the C9orf72 complex strengthens binding to the FIP200 Claw domain. The C9orf72 complex and FLAG-tagged FIP200^Claw^ were phosphorylated separately using substoichiometric amounts of ULK1. ATP was depleted with apyrase, and the indicated combinations were mixed and then incubated with anti-FLAG agarose (for workflow and controls see **Extended Data Fig. 2a**). After washing, bound proteins were eluted and input (I) and elution (E) fractions were analysed by SDS-PAGE and Coomassie staining.

### ULK1-, ULK2- and TBK1-mediated phosphorylation of the C9orf72 complex strengthens its binding to the FIP200 Claw domain

Mirroring its interaction with full-length FIP200 (**Fig. 1c,e**), the C9orf72 complex also bound the isolated FIP200 Claw domain in a manner dependent on ULK1- and TBK1-mediated phosphorylation (**Fig. 2d**). ULK2 is closely related to ULK1 and has largely overlapping, partly redundant functions in canonical autophagy^90^. Consistent with this, ULK2-dependent phosphorylation enhanced binding between the C9orf72 complex and the FIP200 Claw domain to a similar extent as ULK1- and TBK1-dependent phosphorylation, whereas ULK3 had only a modest effect (**Fig. 2d**). Notably, lambda protein phosphatase (λ-PP) treatment strongly reduced binding (**Fig. 2d**) suggesting that at least a subset of the regulatory phosphorylation sites is present on one or both interacting proteins. Because the FIP200 Claw domain was expressed in *E. coli* and is therefore unlikely to be phosphorylated, this suggests that the insect cell-produced C9orf72 complex carries at least some of the relevant regulatory phosphorylation sites that promote binding to the Claw domain. To test this, and to determine whether it is indeed phosphorylation of the C9orf72 complex that drives binding to the FIP200 Claw domain, we phosphorylated either the C9orf72 complex or the FIP200 Claw domain individually using substoichiometric amounts of recombinant ULK1, depleted ATP with apyrase, and then added the corresponding non-phosphorylated binding partner (**Extended Data Fig. 2a**). Strikingly, phosphorylation of the C9orf72 complex, but not of the FIP200 Claw domain, enhanced complex formation (**Fig. 2e**). Together, these findings indicate that kinases regulating bulk and selective autophagy can enhance the C9orf72 complex-FIP200 Claw domain interaction by phosphorylating the C9orf72 complex. On this basis, we used ULK1 and TBK1 interchangeably in subsequent *in vitro* experiments.

### The SMCR8 subunit mediates direct binding of the C9orf72 complex to the FIP200 Claw domain

To better understand how the C9orf72 complex interacts with the FIP200 Claw domain, we purified different C9orf72 complex subunits and subcomplexes and tested their binding to the FIP200 Claw domain. The SMCR8-C9orf72 subcomplex exhibited binding comparable to the intact C9orf72 complex (**Fig. 3a**), suggesting that the WDR41 subunit is dispensable for FIP200 binding. To further dissect which component of the SMCR8-C9orf72 subcomplex mediates this interaction, we attempted to express and purify the individual subunits. As previously reported, SMCR8 could not be purified in isolation^91^. However, the C9orf72 subunit - which could be purified alone after dissociating from the C9orf72-SMCR8 subcomplex during ion-exchange chromatography - completely lost its ability to bind the FIP200 Claw domain (**Fig. 3a**), indicating that SMCR8 mediates the interaction between the C9orf72 complex and the FIP200 Claw domain.

**Fig. 3.**
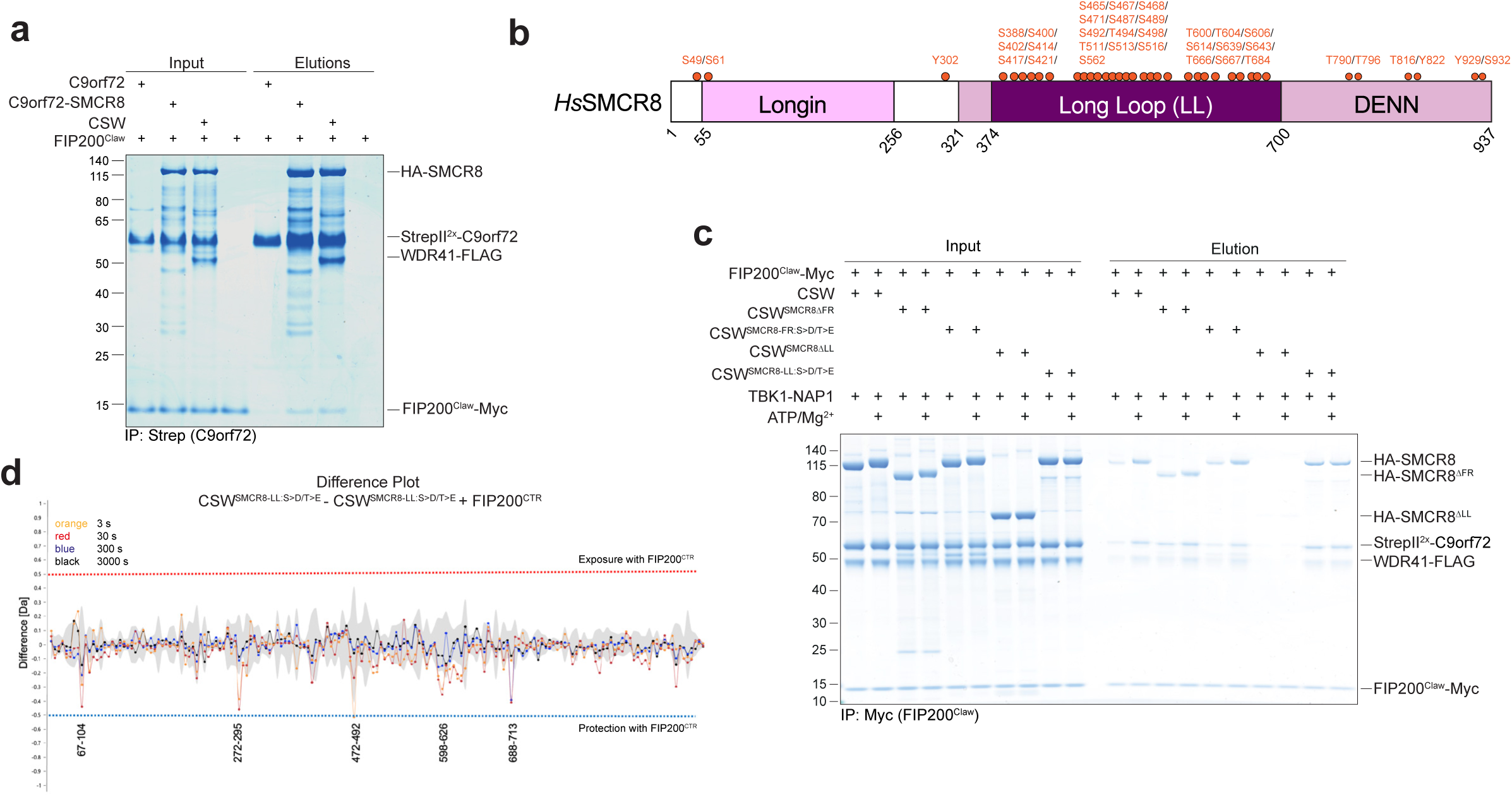
Phosphorylation of the SMCR8 long loop promotes FIP200 Claw domain binding. **a**, SMCR8 mediates binding of the C9orf72 complex to the FIP200 Claw domain. Pulldown assays were performed to compare binding of the FIP200 Claw domain to the intact C9orf72 complex (CSW), the SMCR8-C9orf72 subcomplex, and C9orf72 alone. C9orf72 and the indicated C9orf72-containing complexes were immobilized on Strep-Tactin resin and incubated with the FIP200 Claw domain. After washing, bound proteins were eluted, and input and elution fractions were analyzed by SDS-PAGE followed by Coomassie staining. **b**, Schematic domain overview of *H. sapiens* SMCR8. SMCR8 comprises an N-terminal Longin fold and a C-terminal DENN domain. The DENN domain contains an extended intrinsically disordered insertion, termed the long loop (LL; residues 374–700). Previously reported SMCR8 phosphorylation sites from PhosphoSitePlus database (v6.8.2)^92,97^ are indicated by red dots. **c**, Deletion of the SMCR8 long loop largely abolishes binding of the C9orf72 complex to the FIP200 Claw domain, and TBK1-mediated phosphorylation does not rescue this defect. C9orf72 complexes (wild type or the indicated mutants) were phosphorylated with TBK1-NAP1 (1:20 relative to the C9orf72 complex) or mock-treated by incubation with TBK1-NAP1 under identical conditions without ATP/Mg²⁺. C9orf72 complex mutants carried either deletions of the SMCR8 long loop (CSW^SMCR8ΔLL^) or flexible region (CSW^SMCR8ΔFR^) or phosphomimicking substitutions of all Ser/Thr residues within these regions (CSW^SMCR8-FR:S>D/T>E^ and CSW^SMCR8-LL:S>D/T>E^). CSW complexes were then incubated with Myc-tagged FIP200 Claw domain and captured on Myc resin. After washing, bound proteins were eluted, and input (post-treatment) and elution fractions were analysed by SDS-PAGE followed by Coomassie staining. **d**, HDX-MS analysis of the phosphomimicking SMCR8 long loop mutant in the context of the C9orf72 complex (CSW^SMCR8-LL:S>D/T>E^) upon binding the FIP200^CTR^. Difference plot showing changes in deuterium uptake for the SMCR8^LL:S>D/T>E^ subunit within the CSW^SMCR8-LL:S>D/T>E^ complex in the presence of the FIP200^CTR^. Negative values approaching or exceeding the significance threshold (-0.5) indicate regions protected upon complex formation. Deuterium uptake was measured at four time points: 3 s (orange), 30 s (red), 300 s (blue), and 3000 s (black).

### Phosphorylation of the SMCR8 long loop regulates FIP200 Claw domain binding

SMCR8 adopts an N-terminal Longin fold followed by a C-terminal DENN (differently expressed in normal and neoplastic cells) domain^24^ (**Fig. 3b**). The DENN domain contains an extended intrinsically disordered insertion - hereafter referred to as the long loop - spanning amino acids 374–700 (**Fig. 3b** and **Extended Data Fig. 3a**). SMCR8 harbours numerous reported phosphorylation sites, most of which cluster within the long loop^92^ (**Fig. 3b**). Given that our data indicate that the interaction between the C9orf72 complex and FIP200 is regulated by phosphorylation of the C9orf72 complex, and in light of the phosphorylation-dependent electrophoretic mobility shift of SMCR8 upon ULK1-, ULK2-and TBK1-mediated phosphorylation (**Fig. 1d,e** and **Fig. 2d**), we hypothesised that phosphorylation of the SMCR8 long loop modulates binding to the FIP200 Claw domain.

Consistently, substituting all serine and threonine residues in the SMCR8 long loop with the corresponding phosphomimicking residues aspartate and glutamate (CSW^SMCR8-LL:S>D/T>E^) markedly increased the affinity of the C9orf72 complex for the FIP200 Claw domain (**Fig. 3c**), supporting the idea that phosphorylation within the SMCR8 long loop promotes binding and that phosphomimicking substitution can bypass the requirement for phosphorylation to achieve high-affinity binding. To further test this interaction and define the SMCR8 binding site in greater detail, we used hydrogen-deuterium exchange mass spectrometry (HDX-MS) to compare deuterium incorporation in the phosphomimicking-loop C9orf72 complex mutant (CSW^SMCR8-LL:S>D/T>E^) in the presence and absence of the FIP200^CTR^. Upon complex formation, we observed reduced deuterium incorporation for a subset of SMCR8-derived peptides (**Fig. 3d**). One protected region mapped to the SMCR8 long loop, whereas a second protected region (residues 272–295) lay within another largely disordered segment hereafter termed the flexible region (FR; residues 191-323; **Extended Data Fig. 3a**). Similar to the long loop, the FR is largely unresolved in currently available cryo-EM structures of the C9orf72 complex^27,28,91,93^. However, substituting all serine and threonine residues in this flexible region with phosphomimicking residues (CSW^SMCR8-FR:S>D/T>E^), or deleting the region entirely (CSW^SMCR8ΔFR^), did not impair FIP200 Claw-domain binding (**Fig. 3c**). In contrast, deletion of the SMCR8 long loop (CSW^SMCR8ΔLL^) largely abolished binding of the C9orf72 complex to the FIP200 Claw domain, with TBK1-mediated phosphorylation unable to rescue this defect (**Fig. 3c**). Together, these data establish the SMCR8 long loop as the key C9orf72 complex element that mediates direct binding to the FIP200 Claw domain and demonstrate that phosphorylation within this loop significantly strengthens this interaction.

### Two phosphoregulated FIR motifs in the SMCR8 long loop mediate FIP200 binding

The FIP200 Claw domain has previously been shown to interact with proteins containing FIR (FIP200-interacting region) motifs^57,59–61,94,95^. FIR motifs are short linear sequences with the consensus ψψΘxxΓ, where Θ is typically a hydrophobic or aromatic residue, Γ is usually a bulky hydrophobic residue (most commonly leucine or isoleucine), x denotes any residue, and ψ is either an acidic residue or a serine or threonine (Ser/Thr). Throughout this study, positions within the FIR motif are numbered relative to Θ (Θ = 0), such that the two ψ residues occupy the -2 and -1 positions and Γ is at +3. Phosphorylation within FIR motifs - often involving Ser/Thr residues at the ψ positions - can enhance binding to FIP200, as the resulting negative charge engages a positively charged patch within the FIP200 Claw domain, including lysine 1569 (K1569) and arginine 1573 (R1573)^60^.

Strikingly, inspection of the protected long-loop region identified by HDX-MS (residues 472-492; **Fig. 3d**) revealed a sequence with clear similarity to previously reported FIR motifs (**Fig. 4a**). To test whether the SMCR8-FIP200 interaction is FIR motif-dependent, we examined a Claw-domain mutant (FIP200^Claw-K1569A/R1573E^) previously reported to disrupt FIR motif binding^57,60^. Unlike the wild-type Claw domain, this mutant completely lost the ability to interact with the C9orf72 complex (**Fig. 4b**), indicating that the interaction is mediated by FIR motif(s).

**Fig. 4.**
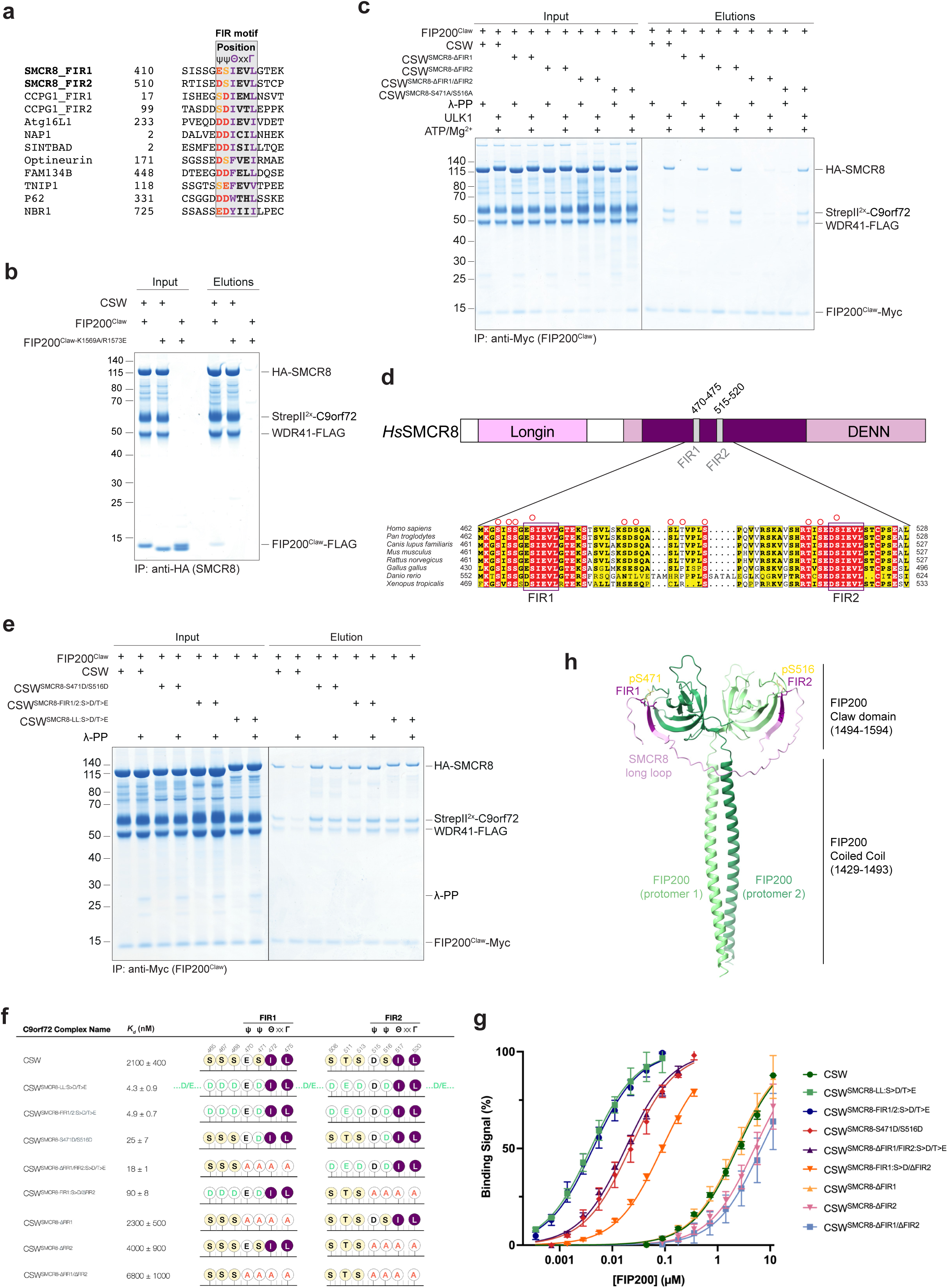
Two phosphoregulated FIR motifs in the SMCR8 long loop mediate high-affinity binding to the FIP200 Claw domain. **a**, Sequence alignment of the two SMCR8 FIR motifs - FIR1 (residues 470-475) and FIR2 (residues 515-520) - with previously reported FIR motifs. The core FIR motif is shaded in grey. The highly conserved residues in the Θ position (typically a hydrophobic or aromatic residue) and Γ position (a bulky hydrophobic residue) are highlighted in purple, while preceding acidic amino acids or serines are marked in red and orange, respectively. **b**, Binding of the C9orf72 complex to the FIP200 Claw domain is abolished by mutation of the FIR-binding site. Pulldown of the HA-tagged C9orf72 complex using a wild-type or FIR-binding defective FIP200 Claw domain (FIP200^Claw-K1569A/R1573E^). **c**, Two FIR motifs in the SMCR8 long loop regulate binding to the FIP200 Claw domain. Recombinant C9orf72 complexes (wild type and indicated mutants) were either dephosphorylated with λ-PP or phosphorylated with recombinant ULK1 at a highly substoichiometric ratio (1:20 relative to the C9orf72 complex). C9orf72 complexes were then incubated with the Myc-tagged FIP200 Claw domain and captured on Myc resin. After washing, bound proteins were eluted, and input (post-treatment) and elution fractions were analysed by SDS-PAGE followed by Coomassie staining. **d**, Domain overview of *H. sapiens* SMCR8, with the two FIR motifs within the long loop indicated (upper panel). Sequence alignment of SMCR8 from multiple organisms highlighting the conservation of the two FIR motifs and reported phosphorylation sites (red circles) within this region (lower panel). **e**, Phosphomimicking of core FIR motif serines is sufficient for Claw binding, with N-terminal phosphorylation sites further enhancing affinity. C9orf72 complexes (wild type and indicated mutants) were either dephosphorylated using λ-PP or left untreated. C9orf72 complexes were then incubated with the Myc-tagged FIP200 Claw domain and immobilized on Myc resin. After washing, bound proteins were eluted, and input and elution fractions were analysed by SDS-PAGE followed by Coomassie staining. **f**, SPR affinity measurements between the extended C-terminal region of FIP200 (aa 1278–1594) and the indicated C9orf72 complex mutants, with mutations in the SMCR8 subunit highlighted in green or red (for phosphomimicking and alanine substitutions, respectively). Equilibrium dissociation constants (*K_d_*) were determined from the SPR sensorgrams (**Extended Data Fig. 4c-k**) by steady-state affinity analysis, as described in the Materials and Methods. All *K_d_* values are based on the dimer concentration of FIP200. *K_d_* values are shown as mean ± standard deviation (SD) from n = 3-7 experiments. **g**, Response-versus-concentration plots from SPR measurements with fitted hyperbolic binding curves. To enable direct comparison of datasets acquired at different surface loadings, steady-state response values were normalised to the maximal response (R_max_) of each dataset, which was set to 100% binding signal. Data points and error bars represent mean ± SD from n = 3-7 experiments. **h**, AF3 model of the C-terminus of FIP200 (residues 1391-1594) in complex with an SMCR8 segment (residues 421-540) that binds the FIP200 Claw domain in a FIR motif-dependent manner. Owing to very low pIDDT scores at the termini of the chosen fragments, only FIP200 residues 1429-1594 and SMCR8 residues 462-526 are shown; S471 and S516 are modelled as phosphorylated. The model includes the FIP200 coiled-coil region preceding the Claw domain. FIP200 dimerises, and the two protomers are shown in different shades of green. SMCR8 is shown in pink, with the experimentally validated FIR motifs highlighted in dark purple and the phosphorylated serines (pS471 and pS516) indicated in light yellow.

However, alanine substitution of the putative SMCR8 FIR motif (hereafter FIR1) in the context of the C9orf72 complex (CSW^SMCR8-ΔFIR1^) caused only a very modest reduction in ULK1/ULK2/TBK1-dependent FIP200 Claw domain binding (**Fig. 4c** and **Extended Data Fig. 4a,b**), suggesting that additional elements within the SMCR8 long loop contribute to the interaction. We therefore reanalysed the long loop and performed a sequence alignment against previously reported FIR motifs (**Fig. 4a**). This analysis identified a second FIR motif (hereafter FIR2) in close proximity (residues 515-520; **Fig. 4d**). Strikingly, alanine substitution of FIR2 alone (CSW^SMCR8-ΔFIR2^) also failed to significantly impair binding to the FIP200 Claw domain (**Fig. 4c** and **Extended Data Fig. 4a,b**), indicating that neither FIR motif is individually required. In contrast, simultaneous disruption of both motifs (CSW^SMCR8-ΔFIR1/ΔFIR2^) strongly impaired both basal and ULK1/ULK2/TBK1-stimulated binding (**Fig. 4c** and **Extended Data Fig. 4a**), establishing FIR1 and FIR2 as functionally redundant FIR motifs that together constitute the principal determinants of Claw domain engagement.

Although the CSW^SMCR8-ΔFIR1/ΔFIR2^ mutant showed markedly reduced binding, a weak phosphorylation-dependent residual interaction was still detectable (**Fig. 4c** and **Extended Data Fig. 4a**). This residual binding may reflect secondary, lower-affinity phosphorylation-stimulated contacts between the C9orf72 complex and the FIP200 Claw domain that are independent of FIR1 and FIR2. Consistent with this possibility, our HDX–MS analyses highlighted additional SMCR8 regions - both within and outside the long loop - that exhibited protection upon FIP200 Claw domain binding (**Fig. 3d**) and could contribute to the weak residual interaction. Notably, however, deletion of the long loop in the C9orf72 complex (CSW^SMCR8ΔLL^) largely abolished binding to the FIP200 Claw domain and could not be rescued by TBK1-mediated phosphorylation (**Fig. 3c**), suggesting that any residual interaction is likely mediated by features within the SMCR8 long loop.

Notably, despite the high sequence variability typical of intrinsically disordered regions, both FIR motifs are highly conserved across species (**Fig. 4d**). Consistent with the FIR consensus motif, FIR1 and FIR2 each contain an acidic residue (E470 and D515, respectively) and an adjacent serine (S471 and S516, respectively) at the ψ positions (**Fig. 4a,d**). Both serines have previously been reported as phosphorylation sites^96,97^ (**Fig. 3b**), providing a plausible molecular mechanism for the TBK1 and ULK1/2-dependent enhancement of binding.

Notably, mutating both core FIR motif serines to alanine (CSW^SMCR8-S471A/S516A^) still allowed ULK1-mediated phosphorylation to stimulate Claw-domain binding to near wild-type levels (**Fig. 4c**). This suggests that, when the core FIR serines are unavailable, phosphorylation at nearby Ser/Thr residues can partially compensate, consistent with the phosphorylation-enhanced FIR-Claw domain interaction reported for p62^57^.

To understand the phosphoregulation in more detail, we generated two phosphomimicking C9orf72 complex mutants and assessed FIP200 Claw domain binding following λ-PP treatment. In the first mutant, the core serine in each FIR motif was replaced with aspartate (CSW^SMCR8-S471D/S516D^). In the second, these sites - together with three upstream Ser/Thr residues in each motif - were substituted with aspartate or glutamate, respectively (CSW^SMCR8-FIR1/2:S>D/T>E^; S465D, S467D, S468D, S471D, S508D, T511E, S513D, S516D). Whereas the wild-type C9orf72 complex (which is phosphorylated during expression in insect cells) showed a strong reduction in FIP200 Claw domain binding after λ-PP treatment, both phosphomimicking mutants retained binding despite dephosphorylation (**Fig. 4e**). Notably, while the extended phosphomimicking mutant, CSW^SMCR8-FIR1/2:S>D/T>E^, bound the Claw domain comparably to the long-loop phosphomimicking variant (CSW^SMCR8-LL:S>D/T>E^), the CSW^SMCR8-S471D/S516D^ showed slightly reduced binding upon dephosphorylation (**Fig. 4e**). These data indicate that phosphomimicking substitution of the core FIR serines is sufficient to promote Claw domain binding, while additional phosphorylation events N-terminal of both core FIR motifs can further enhance affinity. Consistent with this interpretation, S465, S467, S468, S471, T511, S513 and S516 have all been reported as phosphorylation sites *in vivo*^92,97^ (**Fig. 3b**). Finally, ULK1/2- and TBK1-dependent regulation of the SMCR8-FIP200 interaction is consistent with their reported consensus motifs, which favour a hydrophobic residue at the +1 position (often isoleucine) immediately C-terminal to the phosphorylation site^82,98^, with both core FIR motif serines (S471 and S516) and two upstream sites (S465 and T511) matching this motif (**Fig. 4a,d**).

To further probe the functional overlap between the two FIR motifs, we generated C9orf72 complex mutants in which one motif was “primed” for binding via phosphomimicking substitutions at the core serine and three upstream Ser/Thr residues, while the other motif was inactivated by alanine substitution (CSW^SMCR8-FIR1:S>D/ΔFIR2^ and CSW^SMCR8-ΔFIR1/FIR2:S>D/T>E^, respectively). Both constructs retained Claw domain binding after λ-PP treatment (**Extended Data Fig. 4b**), indicating that a single phosphorylated FIR motif is sufficient to support the interaction. Notably, subsequent phosphorylation by ULK1, ULK2, or TBK1 further enhanced binding, suggesting that additional Ser/Thr sites outside the phosphorylated FIR motif can contribute to phosphorylation-dependent strengthening of the SMCR8-FIP200 interaction. Together, these data indicate that the two FIR motifs exhibit functional overlap, while maximal binding is achieved when both motifs are available for engagement and phosphorylation-dependent reinforcement.

To complement the pulldown assays and quantify binding affinities between the C9orf72 complex and FIP200, we performed surface plasmon resonance (SPR) experiments (**Fig. 4f,g** and **Extended Data Fig. 4c-k**). Consistent with our pulldown data (**Fig. 4c,e** and **Extended Data Fig. 4b**), the dephosphorylated wild-type complex bound only weakly to the FIP200^CTR^, with a dissociation constant (*K_d_*) of 2.1 µM (**Fig. 4f,g** and **Extended Data Fig. 4c**). Alanine substitutions in both FIR motifs (CSW^SMCR8-ΔFIR1/ΔFIR2^) reduced affinity ∼3-fold (*K_d_* = 6.8 µM), whereas the individual FIR mutants (CSW^SMCR8-ΔFIR1^ and CSW^SMCR8-ΔFIR2^) showed intermediate affinities (*K_d_* = 2.3 µM and 4.0 µM, respectively; **Fig. 4f,g** and **Extended Data Fig. 4d-f**), indicating that FIR2 contributes slightly more to basal binding in the dephosphorylated state.

Phosphomimicking the entire SMCR8 long loop (CSW^SMCR8-LL:S>D/T>E^) increased affinity by roughly 500-fold (*K_d_* = 4.3 nM; **Fig. 4f,g** and **Extended Data Fig. 4g**). When phosphomimicking substitutions were restricted to the core FIR serines alone (CSW^SMCR8-S471D/S516D^; *K_d_* = 25 nM; **Fig. 4f,g** and **Extended Data Fig. 4h**), affinity remained markedly enhanced relative to the dephosphorylated wild-type complex, but was weaker than that of a mutant additionally carrying phosphomimicking substitutions at the three upstream Ser/Thr residues (CSW^SMCR8-FIR1/2:S>D/T>E^; *K_d_* = 4.9 nM; **Fig. 4f,g** and **Extended Data Fig. 4i**), indicating that phosphomimicking the core FIR serines is sufficient to drive a substantial affinity increase, whereas phosphomimicking the upstream Ser/Thr residues can further strengthen binding.

Because FIP200 forms dimers with the two C-terminal Claw domains in close proximity^57,60^, the presence of two FIR motifs within the SMCR8 long loop - separated by 39 amino acids, a spacing that is largely conserved and never shorter across all analysed organisms (**Fig. 4d**) - suggested a bivalent binding mechanism in which a single C9orf72 complex can engage both Claw domains simultaneously. Consistent with this idea, in our AlphaFold3 (AF3) prediction^99^ each Claw domain is occupied by one of the two FIR motifs, with each motif adopting a binding mode similar to that of previously characterised FIR motif-Claw domain interactions (**Fig. 4h** and **Extended Data Fig. 4l-q**).

To determine whether two FIR motifs are required for the highest-affinity interaction, we deleted one FIR motif and introduced phosphomimicking substitutions at the four Ser/Thr residues associated with the remaining motif. In both cases, affinity was markedly reduced relative to the mutant with both FIR motifs phosphomimicked (CSW^SMCR8-FIR1/2:S>D/T>E^), with a *K_d_* = 18 nM upon deletion of FIR1 (CSW^SMCR8-ΔFIR1/FIR2:S>D/T>E^) and a *K_d_* = 90 nM upon deletion of FIR2 (CSW^SMCR8-FIR1:S>D/ΔFIR2^), corresponding to ∼3-fold and ∼20-fold decreases, respectively (**Fig. 4f,g** and **Extended Data Fig. 4j,k**). The more pronounced affinity decrease upon FIR2 deletion is consistent with its greater contribution to basal binding as observed by SPR.

Together, these results support a model in which the exceptionally high affinity of the phosphorylated C9orf72 complex arises from avidity-driven, bivalent engagement of the two FIP200 Claw domains by FIR1 and FIR2. Collectively, our data establish FIR1 and FIR2 as *bona fide* phosphoregulated motifs within SMCR8 whose phosphorylation by ULK1/2 and TBK1 promotes robust association of the C9orf72 complex with the ULK1/2 complex.

Due to their similarity to previously described non-canonical LC3-interacting region (LIR) motifs and the comparable principle of phosphorylation-dependent regulation reported for these motifs^61,100–103^, we asked whether the SMCR8 FIR motifs might also function as non-canonical LIRs. We therefore tested whether they could mediate recruitment of the C9orf72 complex to the expanding phagophore through interactions with LC3- or GABARAP-family proteins.

Because LIR-mediated binding is often enhanced by phosphorylation of Ser/Thr residues immediately N-terminal to the motif, we tested the phosphomimicking C9orf72 complex mutant in which the core FIR motif serines and the three upstream Ser/Thr residues were substituted with aspartate or glutamate (CSW^SMCR8-FIR1/2:S>D/T>E^). We then tested binding to a panel of LC3 and GABARAP family proteins. Under these conditions, the C9orf72 complex showed no detectable interaction with any LC3- or GABARAP-family member (**Extended Data Fig. 4r**), indicating that the SMCR8 FIR motifs do not function as LIR motifs and are unlikely to mediate direct recruitment of the C9orf72 complex to autophagosomal membranes via LC3- or GABARAP-family proteins.

### Phosphorylation of SMCR8 FIR motifs, but not S402/T796, regulates the C9orf72 complex-ULK1/2 complex interaction in cells

To assess the physiological relevance of our findings, we tested the purified recombinant wild-type C9orf72 complex and variants harbouring mutations in SMCR8 for their ability to co-immunoprecipitate FIP200 and other ULK1/2 complex subunits from HEK293 cell lysates. The CSW^SMCR8-S471A/S516A^ variant, in which the two core FIR-motif serines were replaced with alanine, showed markedly reduced binding to FIP200 and to all other ULK1/2 subunits compared with the wild-type complex (**Fig. 5a**). This defect was mildly exacerbated when the remaining core FIR-motif residues were also substituted with alanine (CSW^SMCR8-ΔFIR1/ΔFIR2^; **Fig. 5a**), further supporting that phosphorylation of the core FIR motif serines is a key determinant of the C9orf72 complex ULK1/2 complex interaction.

**Fig. 5.**
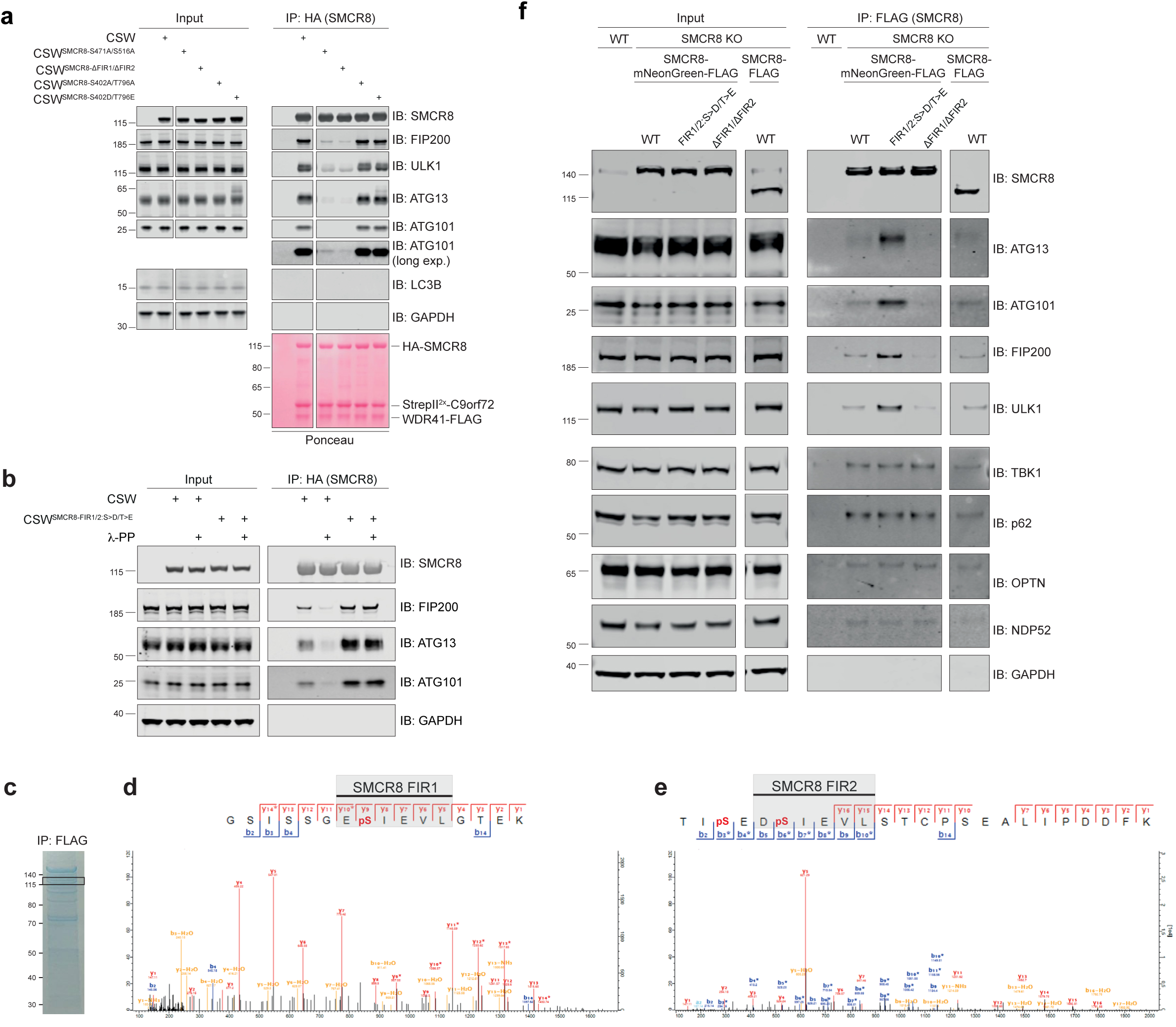
The C9orf72 complex interaction with the ULK1/2 complex depends on the two SMCR8 FIR motifs and their phosphorylation in cells. **a**, Co-immunoprecipitation of FIP200 and ULK1/2 complex subunits by recombinant C9orf72 complex variants. *FIP200*-KO HEK293T cells were rescued with StrepII^2x^-FIP200-FLAG, and lysates were divided into equal aliquots before incubation with purified recombinant C9orf72 complexes containing HA-tagged SMCR8 variants (WT, S471A/S516A, ΔFIR1/ΔFIR2, S402A/T796A or S402D/T796E). Complexes were captured by anti-HA immunoprecipitation (IP). Input and IP fractions were immunoblotted for the indicated proteins. Ponceau staining was used to assess recovery of the purified C9orf72 complex variants across IP fractions. A no-bait control was included by omitting the purified C9orf72 complex. **b**, FIR-motif phosphomimicking strengthens C9orf72 complex binding to the ULK1/2 complex. Co-immunoprecipitation was performed as in Fig. 5a, comparing untreated versus λ-PP-treated recombinant wild-type C9orf72 complex and the CSW^SMCR8-FIR1/2:S>D/T>E^ mutant. Input and IP fractions were immunoblotted for the indicated proteins, and a no-bait control was included. **c**, SMCR8 purification from HEK293T cells for mass spectrometry analysis. HEK293T *SMCR8*-KO cells were rescued with doxycycline-inducible StrepII^2x^-SMCR8-FLAG and treated with 1 µg/mL doxycycline for 24 hours. Cells were lysed and subjected to anti-FLAG immunoprecipitation, followed by elution with 3X FLAG peptide. The eluate was analysed by SDS-PAGE and Coomassie staining. **d,e**, Mass spectrometry analysis of FLAG-tagged SMCR8 isolated from HEK293T cells. The boxed region shown in Fig. 5c was excised and analysed by mass spectrometry. Fragment ion spectra for peptides containing the phosphorylated FIR1 (pS471) and FIR2 (pS513 and pS516) are shown in (**d**) and (**e**), respectively. The data were acquired using ion-trap mass spectrometry (ITMS) with higher-energy collisional dissociation (HCD). Assigned b- and y-ions are indicated in blue and red, respectively. **f**, Co-immunoprecipitation of ULK1/2 complex subunits, TBK1 and autophagy cargo adaptors with SMCR8 variants expressed in *SMCR8*-KO HEK293 cells rescued with C-terminal mNeonGreen-FLAG-tagged SMCR8 (wild type, SMCR8^FIR1/2:S>D/T>E^ or SMCR8^ΔFIR1/ΔFIR2^) or with wild-type SMCR8-FLAG. Lysates were subjected to anti-FLAG immunoprecipitation using anti-FLAG agarose, and input and IP fractions were immunoblotted for the indicated proteins. Wild-type HEK293 cells were included as a no-bait control.

To further assess the physiological relevance of the observed phosphoregulation, we tested the recombinant C9orf72 complex variant in which the regulatory Ser/Thr residues across both FIR motifs were replaced with phosphomimicking substitutions (CSW^SMCR8-FIR1/2:S>D/T>E^). Consistent with our model, this mutant co-immunoprecipitated markedly more FIP200, ATG13 and ATG101 from HEK293 lysates than the wild-type complex (**Fig. 5b**). In line with this, λ-PP treatment of the recombinant wild-type complex strongly reduced association with FIP200, ATG13 and ATG101 - presumably by removing regulatory phosphorylations acquired during expression in insect cells - whereas the phosphomimicking CSW^SMCR8-FIR1/2:S>D/T>E^ complex was unaffected by λ-PP treatment (**Fig. 5b**).

Having established that FIR-motif phosphorylation controls ULK1/2 complex engagement, we next asked whether other previously reported TBK1-dependent phosphorylation sites in SMCR8 modulate this interaction. A previous study identified SMCR8 as a TBK1 substrate *in vitro*, and phosphorylation at S402 and T796 was implicated in clearance of cytosolic p62 aggregates^17^. We therefore examined whether phosphomimicking or alanine substitutions at these sites affect binding to the ULK1/2 complex. Both the recombinant CSW^SMCR8-S402D/T796E^ and CSW^SMCR8-S402A/T796A^ complexes co-immunoprecipitated ULK1/2 complex subunits from cell lysates to a similar extent as the wild-type complex (**Fig. 5a**). This is consistent with our *in vitro* assays showing that these substitutions do not measurably affect binding of the corresponding C9orf72 complex variants to the FIP200 Claw domain (**Extended Data Fig. 5a**), supporting the idea that phosphorylation at S402 and T796 influences p62 clearance via a mechanism distinct from ULK1/2 complex recruitment.

To extend these findings beyond *in vitro* and lysate reconstitution and to further probe whether FIR motif phosphorylation also governs ULK1/2 complex engagement in a cellular context, we next examined this interaction in cells. To confirm that the SMCR8 FIR motifs are phosphorylated in cells, we immunoprecipitated FLAG-tagged SMCR8 from HEK293T cells (**Fig. 5c**) and analysed it by mass spectrometry. Consistent with our phosphoregulation model, MS/MS spectra confirmed phosphorylation within both FIR1 and FIR2 (**Fig. 5d,e**).

Next, we generated *SMCR8*-knockout (KO) Flp-In HEK293 cells (**Extended Data Fig. 5b,c**), and reintroduced either wild-type SMCR8 or the two previously characterised mutants SMCR8^FIR1/2:S>D/T>E^ and SMCR8^ΔFIR1/ΔFIR2^ into the genomic safe-harbour locus by Flp-mediated recombination. Each SMCR8 variant carried a C-terminal mNeonGreen-FLAG tag, enabling immunoprecipitation of the corresponding C9orf72 complexes and FACS-based enrichment of cell populations with comparable SMCR8 expression levels for subsequent autophagy flux analyses. Importantly, the mNeonGreen-FLAG tag did not interfere with the phosphorylation-dependent interaction between SMCR8 and the FIP200 Claw domain, as assessed by *in vitro* pulldown experiments (**Extended Data Fig. 5d**), nor with SMCR8-mediated engagement of the ULK1/2 complex in HEK293 cells (**Fig. 5f**). Consistent with our *in vitro* and cell lysate findings, immunoprecipitation of exogenous SMCR8^FIR1/2:S>D/T>E^ recovered substantially higher levels of FIP200 and other ULK1/2 complex subunits, whereas alanine substitution of both FIR motifs (SMCR8^ΔFIR1/ΔFIR2^) further weakened the interaction relative to wild-type SMCR8 (**Fig. 5f**). Together, these results confirm that the C9orf72 complex ULK1/2 complex interaction is FIR motif dependent and regulated by phosphorylation in cells.

The C9orf72 complex interacts not only with the ULK1/2 complex but has also been suggested to associate with TBK1 and the autophagy cargo adaptors p62 and OPTN^17,76^. Surprisingly, although ULK1/2 complex binding was markedly enhanced by the phosphomimicking SMCR8^FIR1/2:S>D/T>E^ mutant, association with TBK1 and the cargo adaptors p62, OPTN and NDP52 was unchanged relative to wild-type SMCR8 or SMCR8^ΔFIR1/ΔFIR2^ (**Fig. 5f**). These data indicate that recruitment of TBK1 and cargo adaptors to the C9orf72 complex is independent of ULK1/2 association, and is likely indirect, consistent with the absence of a detectable direct interaction in our *in vitro* pulldown assays (**Fig. 1c**).

Taken together, these data support that the C9orf72 complex-ULK1/2 complex interaction is mediated by phosphorylation of the SMCR8 FIR motifs, whereas phosphorylation at the TBK1 sites S402 and T796 likely regulates p62 clearance through a mechanism distinct from ULK1/2 complex recruitment.

### The C9orf72 complex-ULK1/2 complex interaction differentially regulates selective autophagy pathways

Having established how the C9orf72 complex engages the ULK1/2 complex, we next sought to define the physiological relevance of this interaction. To this end, we used Flp-In HEK293 cell lines expressing either SMCR8^ΔFIR1/ΔFIR2^ or SMCR8^FIR1/2:S>D/T>E^. We refer to these SMCR8 variants as separation-of-function mutants: SMCR8^ΔFIR1/ΔFIR2^ markedly weakens the SMCR8-FIP200 interaction, whereas SMCR8^FIR1/2:S>D/T>E^ stabilises it and renders it insensitive to phosphorylation by introducing phosphomimicking substitutions at the regulatory Ser/Thr sites. Importantly, these mutants selectively modulate the interaction with FIP200 while leaving other C9orf72 complex functions intact. This is important because the C9orf72 complex also regulates pathways that impinge on autophagy, including lysosomal amino-acid sensing, which modulates mTORC1 activity and thereby regulates TFEB/TFE3-dependent lysosome biogenesis, small GTPase-dependent membrane trafficking and actin-linked vesicle transport^17,19,22,25,27,29,30,104^. Thus, these separation-of-function mutants provide a tool to interrogate autophagy without the pleiotropic effects typically associated with *SMCR8* or *C9orf72* KO or knockdown strategies, allowing direct assessment of how the C9orf72 complex-ULK1/2 complex interaction contributes to autophagy in cells.

To monitor autophagy-dependent lysosomal delivery, we used monomeric Keima (mKeima), a pH-sensitive fluorescent protein whose excitation spectrum is bimodal and shifts with pH: it is preferentially excited at ∼440 nm at neutral pH and at ∼586 nm in acidic environments such as lysosomes, while its emission remains near 620 nm independent of pH. Because mKeima is relatively resistant to lysosomal proteolysis, the ratio of mKeima fluorescence upon excitation at 586 versus 440 nm (emission collected at ∼620 nm) provides a robust readout of lysosomal delivery and is widely used to quantify bulk and selective autophagy (**Fig. 6a,b**)^105,106^.

**Fig. 6.**
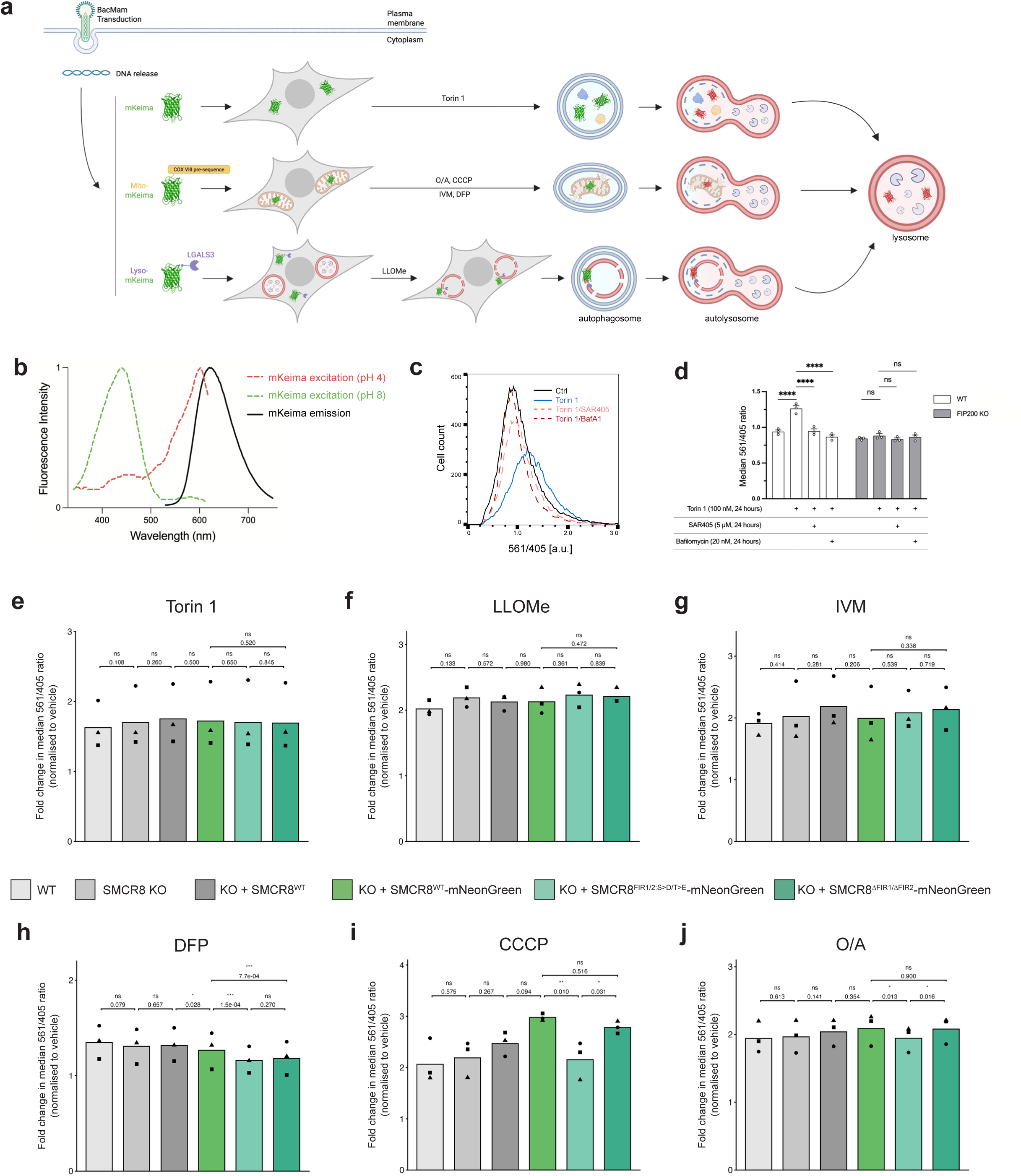
The SMCR8-FIP200 interaction differentially regulates selective autophagy pathways. **a**, Schematic overview of the mKeima-based autophagy flux assay. BacMam transduction is used to transiently express the indicated autophagy reporters in HEK293 cells. The bulk autophagy reporter (mKeima) is cytosolic, whereas the mitophagy reporter (mito-mKeima) is targeted to the mitochondrial matrix via an N-terminal COX VIII pre-sequence. The lysophagy reporter (mKeima-LGALS3) is initially diffuse in the cytosol but, following lysosomal membrane damage, accumulates on the cytosolic surface of ruptured lysosomes. Upon induction of bulk or selective autophagy pathways, the corresponding reporters are sequestered into autophagosomes, which subsequently fuse with lysosomes to form autolysosomes. mKeima is resistant to lysosomal proteolysis, and acidification within autolysosomes shifts its excitation maximum to ∼586 nm; thus, the acidic-to-neutral mKeima fluorescence ratio provides a readout of autophagy flux. **b**, pH-dependent excitation and emission spectra of mKeima. mKeima displays a bimodal excitation spectrum, with a peak at ∼440 nm under neutral pH conditions (green dashed line) and a peak at ∼586 nm under acidic pH conditions (red dashed line). In both cases, the emission maximum is ∼620 nm (solid black line). **c,d**, Autophagic flux analysis by flow cytometry. Wild-type (WT) and *FIP200*-KO HEK293 cells were transduced with BacMam encoding cytosolic mKeima to monitor bulk autophagy. Cells were treated with Torin 1 alone or in combination with SAR405 or bafilomycin A1 (BafA1), as indicated. For each cell, the mKeima excitation ratio was calculated as the 615 nm emission intensity upon 561 nm excitation divided by that upon 405 nm excitation (561/405). (**c**) Representative overlaid histograms of single-cell 561/405 ratios for WT cells under the indicated treatments. (**d**) Quantification of autophagic flux in WT and *FIP200*-KO HEK293 cells shown as the median 561/405 ratio for each sample and bars represent the mean ± SEM of triplicates from a representative experiment. Statistical significance was assessed using a two-way ANOVA followed by Tukey’s multiple-comparisons test. *P* < 0.05 (*), *P* < 0.01 (**), *P* < 0.001 (***), and *P* < 0.0001 (****). **e-j**, Role of the SMCR8-FIP200 interaction in bulk and selective autophagy. Autophagic flux was quantified by flow cytometry in wild-type, *SMCR8*-KO, SMCR8 WT-rescue, and SMCR8 mutant cell lines with decreased or increased interaction with FIP200 (SMCR8^ΔFIR1/ΔFIR2^ and SMCR8^FIR1/2:S>D/T>E^, respectively; cell-line colour key shown below panels **e-g**). For each condition, the treatment response is shown as the fold change in the median mKeima 561/405 excitation ratio relative to the corresponding untreated control. The underlying raw median 561/405 ratios for untreated and treated samples are shown in **Extended Data Fig. 6c–h.** (**e**) Bulk autophagy, induced with 100 nM Torin-1 for 24 h. (**f**) Lysophagy, induced with 1 mM LLOMe for 1 h, followed by an 8 h recovery. (**g–j**) Mitophagy, induced with 16 μM ivermectin (IVM) for 24 h (**g**), 1 mM deferiprone (DFP) for 48 h (**h**), 5 μM CCCP for 6 h (**i**), or 10 μM oligomycin/antimycin A (O/A) for 6 h (**j**). Bars show mean ± s.e.m. (n = 3 independent experiments; dots, experiments; each experiment in biological triplicate). Cell lines expressing mNeonGreen-tagged SMCR8 were plotted alongside cell lines lacking mNeonGreen expression; however, statistical analyses were restricted to the specified comparisons indicated in the figures. Two-sided Wald tests were performed, with *P* values adjusted using Holm’s method to account for multiple comparisons and control the family-wise error rate. *P* < 0.05 (*)*, P < 0.01 (**)*, *P* < 0.001 (***), *P* < 0.0001 (****).

To measure bulk autophagy, we transiently expressed mKeima in our cell lines using BacMam, a non-integrating baculovirus-based gene delivery system for mammalian cells^107,108^ (**Fig. 6a**). Importantly, BacMam transduction of HEK293 cells did not affect autophagy (**Extended Data Fig. 6a**) and was hence used throughout this study. We first validated the assay in wild-type and *FIP200*-KO HEK293 cells by monitoring bulk autophagic flux. To confirm that the increase in flux upon treatment with the bulk autophagy inducer Torin 1 was autophagy-dependent, we carried out the experiments in the presence of two commonly used inhibitors - the VPS34/PIK3C3 inhibitor SAR405 and the lysosomal acidification inhibitor bafilomycin A1 (BafA1) - both of which abolished Torin 1-induced autophagy flux (**Fig. 6c,d**).

To test whether C9orf72 complex-ULK1/2 complex engagement influences autophagy, we next measured bulk autophagy flux in cells expressing the two separation-of-function mutants, SMCR8^ΔFIR1/ΔFIR2^ and SMCR8^FIR1/2:S>D/T>E^ and compared them with wild-type SMCR8. To ensure comparable SMCR8 expression across cell lines, we first FACS-sorted populations expressing the corresponding mNeonGreen-tagged wild-type or mutants SMCR8 protein prior to flux measurements (**Extended Data Fig. 6b**). Neither separation-of-function mutant altered bulk autophagy (**Fig. 6e** and **Extended Data Fig. 6c**), suggesting that the SMCR8-FIP200 interaction is dispensable for bulk autophagy regulation under these conditions.

Next, we examined selective autophagy of lysosomes (lysophagy) and mitochondria (mitophagy), as defects in organelle quality control - particularly impaired clearance of damaged lysosomes and mitochondria - have been implicated in the pathogenesis of neurodegenerative diseases, including ALS and FTD^109,110^.

We first assayed lysophagy using the mKeima-LGALS3 reporter^111^. Neither weakening nor stabilising the SMCR8-FIP200 interaction impaired L-leucyl-L-leucine methyl ester (LLOMe)-induced lysophagy (**Fig. 6f** and **Extended Data Fig. 6d**), indicating that the SMCR8-FIP200 interaction does not regulate LLOMe-induced lysophagy.

We next quantified mitophagy using an mKeima-based reporter targeted to the mitochondrial matrix (mito-mKeima). Mitophagy can proceed via mechanistically distinct routes, including PINK1-Parkin-dependent ubiquitin-driven pathways, receptor-mediated pathways involving receptors such as BNIP3, NIX and FUNDC1, and other Parkin-independent ubiquitin/adaptor-driven pathways^112^. Because different stimuli differentially engage these pathways, we applied a panel of inducers to probe both Parkin-dependent and -independent mitophagy.

We first assessed mitophagy induced by ivermectin (IVM)^113^. Neither *SMCR8*-KO nor expression of the separation-of-function mutants altered IVM-induced mitophagy (**Fig. 6g** and **Extended Data Fig. 6e**), suggesting that the SMCR8-FIP200 interaction is dispensable for this Parkin-independent pathway. Next, we assayed a second Parkin-independent mitophagy pathway using the iron chelator deferiprone (DFP) as an inducer^114^. Strikingly, both separation-of-function mutants displayed a notable defect in DFP-induced mitophagy (**Fig. 6h** and **Extended Data Fig. 6f**).

Finally, to assess Parkin-dependent mitophagy, we induced mitochondrial depolarisation using carbonyl cyanide m-chlorophenylhydrazone (CCCP) or oligomycin/antimycin A (O/A) and transiently co-expressed Parkin together with mito-mKeima. Under both CCCP and O/A treatment, *SMCR8*-KO had no detectable effect on mitophagy (**Fig. 6i,j** and **Extended Data Fig. 6g,h**). By contrast, stabilising the interaction between the C9orf72 complex and the ULK1/2 complex (SMCR8^FIR1/2:S>D/T>E^) reduced mitophagy flux, whereas weakening this interaction (SMCR8^ΔFIR1/ΔFIR2^) had no significant effect (**Fig. 6i,j** and **Extended Data Fig. 6g,h**).

Taken together, these findings suggest that the C9orf72 complex-ULK1/2 complex interaction can selectively and differentially influence distinct branches of selective autophagy, with effects that depend on both the pathway and the inducing stimulus. Importantly, by selectively weakening the C9orf72 complex-ULK1/2 complex interaction, the SMCR8^ΔFIR1/ΔFIR2^ separation-of-function mutant provides a means to disentangle autophagy-specific consequences of impaired C9orf72 complex-ULK1/2 engagement from other pleiotropic defects that may arise when C9orf72 complex levels are reduced, as reported in ALS/FTD associated with C9orf72 repeat expansions. Over time, even modest impairments in selective autophagy could lead to a gradual accumulation of damaged organelles and disease-associated protein aggregates, with age-related declines in cellular quality-control capacity potentially contributing to the typically late-onset nature of ALS/FTD.

## Discussion

*C9ORF72* hexanucleotide repeat expansions are the most common genetic cause of familial ALS/FTD and frequently reduce C9orf72 protein abundance and, as a result, C9orf72 complex levels. Yet how repeat-expansion-associated reduction in C9orf72 complex abundance contributes to autophagy-lysosome dysfunction and previously reported mitochondrial quality-control defects in C9ORF72-ALS/FTD remains unclear. Here we identify a direct, phosphorylation-regulated molecular link between the C9orf72 complex and autophagy initiation: the SMCR8 subunit engages the C-terminal Claw domain of the FIP200 subunit of the ULK1/2 complex via two highly conserved FIR motifs embedded in an extended, intrinsically disordered loop within SMCR8. Phosphorylation by ULK1/2 or TBK1 strongly potentiates this interaction, enabling avidity-driven, high-affinity binding through simultaneous engagement of the two Claw domains in the FIP200 dimer. In cells, SMCR8 separation-of-function mutants that selectively weaken or stabilise the SMCR8-FIP200 interaction tune ULK1/2 complex association and reveal stimulus-dependent, pathway-selective effects on mitophagy, while leaving bulk autophagy and lysophagy largely intact. Together, these findings define a regulated C9orf72-ULK1/2 complex axis and provide a mechanistic framework by which repeat-expansion-associated reduction in C9orf72 complex abundance may contribute to previously reported defects in mitochondrial quality control in C9ORF72-ALS/FTD.

FIP200 is increasingly recognised as a central recruitment hub in autophagy, integrating inputs from cargo adaptors, regulatory factors and core autophagy machinery^33,36,115,116^. FIR motif-dependent FIP200 Claw domain recognition has emerged as a common mechanism by which adaptors recruit the ULK1/2 complex to selective autophagy cargos to promote cargo-proximal autophagosome formation. In several cases, phosphorylation within FIR motifs or at adjacent Ser/Thr residues strengthens Claw engagement^57–60^, providing a means to tune the strength or duration of ULK1/2 complex recruitment. Our work expands this framework by identifying SMCR8 as a phosphorylation-regulated Claw domain interactor within the ALS/FTD-linked C9orf72 complex, placing SMCR8 alongside other FIR-containing autophagy proteins that engage FIP200. SMCR8 has not been established as a canonical cargo receptor, and its FIR motifs do not detectably bind LC3/GABARAP-family proteins, supporting the idea that SMCR8’s FIR module is specialised for regulated ULK1/2 complex binding rather than cargo-directed ULK1/2 complex recruitment and direct phagophore tethering.

A notable feature of the SMCR8-FIP200 interaction is the presence of two conserved FIR motifs within the SMCR8 long loop. FIR1 and FIR2 are individually dispensable yet together support maximal binding, and phosphomimicking substitutions drive low-nanomolar binding affinities. Consistent with a contribution from multivalency, removing one motif while priming the other reduces affinity by several-fold (**Fig. 4f,g**), indicating that the second motif measurably enhances overall engagement - an effect compatible with avidity within the dimeric FIP200 scaffold. Moreover, the conserved spacing between FIR1 and FIR2 is consistent with engagement of both Claw domains, and our AF3 model (**Fig. 4h**) places the two FIR motifs on separate Claw domains. This tandem-motif arrangement provides a plausible explanation for the pronounced phosphorylation sensitivity of complex formation: incremental increases in negative charge distributed across both FIR motifs can cooperatively stabilise SMCR8-FIP200 association, thereby allowing small increases in phosphorylation to produce a strong increase in binding without requiring global changes in ULK1/2 activity.

Our findings further place the C9orf72 complex within an autophagy-relevant kinase network by showing that both ULK1/2 and TBK1 can potentiate SMCR8-FIP200 binding. ULK1/2 are activated during autophagy initiation and phosphorylate multiple proteins that coordinate early autophagosome biogenesis^46,117,118^, whereas TBK1 is a central regulator of selective autophagy and is genetically linked to ALS/FTD^119–121^. The ability of these kinases to strengthen the SMCR8-FIP200 interaction suggests a point of convergence at which phosphorylation inputs linked to selective autophagy, particularly mitophagy, can tune the likelihood and stability of C9orf72 complex-ULK1/2 complex association. This is consistent with a local reinforcement model in which kinase activity at nascent initiation sites stabilises C9orf72 complex engagement with FIP200 in a stimulus- and site-specific manner. While phosphorylation is increasingly recognised to modulate FIR-Claw interactions, the upstream kinase(s) have been defined for only a subset of interactors (e.g., TBK1 for optineurin and TNIP1^59,60^), and remain unclear for many others. Our results suggest that ULK1 and ULK2 can act upstream of a FIR motif-mediated interaction to directly promote FIP200 Claw-domain engagement by phosphorylating the FIR-containing interactor itself.

Notably, FIR-motif phosphorylation does not alter C9orf72 complex association with TBK1 or with the tested cargo adaptors under our conditions (**Fig. 5f**). Consistent with this, our *in vitro* pulldown assays did not detect direct interactions between the C9orf72 complex and TBK1-NAP1 or the cargo adaptor OPTN (**Fig. 1c**). Together, these observations suggest that regulation of C9orf72 complex-ULK1/2 complex engagement via the SMCR8 FIR motifs is mechanistically separable from reported C9orf72 complex associations with TBK1 and cargo adaptors, which may instead reflect indirect interactions in cells.

A persistent challenge in the C9orf72 complex field has been the heterogeneity of autophagy-related phenotypes reported upon C9orf72 or SMCR8 knockdown/knockout across different models, including both impaired and enhanced autophagy-associated readouts^17,19,20,122,123^. This divergence likely reflects differences in cell type, perturbation regime and assay readouts, together with the pleiotropic roles of the C9orf72 complex beyond autophagy. In addition to associating with the autophagy initiation machinery, the C9orf72 complex participates in membrane trafficking and lysosomal nutrient-sensing pathways that influence mTORC1 activity and TFEB/TFE3-dependent transcriptional programs^22,29,30,104,124^. SMCR8 depletion can therefore confound autophagy readouts through altered lysosome homeostasis, membrane trafficking, transcriptional states and compensatory rewiring. To isolate the specific contribution of C9orf72 complex recruitment to the autophagy initiation machinery from these broader effects, we used SMCR8 separation-of-function mutants that selectively weaken or stabilise the SMCR8-FIP200 interaction while preserving overall C9orf72 complex integrity. This bidirectional interaction-tuning strategy, combined with quantitative autophagy flux reporters, enabled direct, internally consistent comparisons across bulk and selective autophagy pathways.

Using this approach, we observed that SMCR8-FIP200 coupling exerts stimulus-dependent, pathway-selective effects on mitophagy, while having minimal impact on bulk autophagy or lysophagy under our conditions. Neither weakening nor stabilising SMCR8-FIP200 binding measurably alters bulk autophagy flux, nor does it affect LLOMe-induced lysophagy or ivermectin-induced mitophagy. In contrast, stabilising the interaction suppresses CCCP and O/A-induced Parkin-dependent mitophagy, while both stabilisation and weakening impair DFP-induced mitophagy. These distinct response patterns argue for pathway-specific control logic and suggest that some mitophagy routes are especially sensitive to the extent and dynamics of C9orf72 complex-ULK1/2 complex engagement. In Parkin-driven settings, weakening SMCR8-FIP200 coupling is buffered in our assay, whereas constitutive stabilisation suppresses mitophagy. This is consistent with the possibility that excessive or prolonged SMCR8-FIP200 engagement becomes non-productive in Parkin-driven mitophagy, without detectably altering cargo adaptor/TBK1 association with FIP200 (**Fig. 5f**). Thus, the C9orf72 complex-FIP200 interaction does not appear to be rate-limiting under these conditions, but nor is it functionally neutral: weakening is buffered, whereas forced stabilisation reveals a requirement for dynamic turnover. More broadly, our separation-of-function phenotypes support the idea that the phosphorylation-dependent C9orf72 complex-FIP200 interaction must remain reversible: dynamic cycles of phosphorylation and dephosphorylation - driven by local ULK1/2 and TBK1 activity and opposed by autophagy-linked phosphatases^125–127^ - may help prevent persistent C9orf72 complex accumulation and maintain dynamic exchange of FIR-motif-containing factors during autophagy initiation. By contrast, DFP-induced mitophagy may depend on a Goldilocks level of C9orf72 complex-ULK1/2 complex association, in which both insufficient and excessive engagement become non-productive either by failing to support productive initiation or by stabilising the SMCR8-FIP200 interaction in a locked state. These results underscore that distinct mitophagy triggers can engage partially separable regulatory inputs even when converging on common core machinery.

Finally, our findings offer a concrete mechanistic hypothesis for how repeat-expansion-associated reduction in C9orf72 complex abundance could contribute to selective cellular vulnerabilities in C9ORF72-ALS/FTD. Mitophagy is a central pillar of mitochondrial quality control, and mitochondrial quality-control defects have been repeatedly implicated in ALS/FTD^77–79^. Even without directly titrating C9orf72 complex abundance, our data identify a phosphorylation-tunable SMCR8-FIP200 interface that may be particularly sensitive to reduced C9orf72 complex levels and thereby selectively impact mitophagy. Importantly, this same interface is potentiated by TBK1, a kinase genetically linked to ALS/FTD, suggesting a convergent vulnerability: reduced C9orf72 complex abundance could be compounded by compromised TBK1-dependent phosphorylation, further narrowing the conditions under which productive ULK1/2 complex engagement can be achieved. Our data do not address the relative contribution of this loss-of-function mechanism compared with repeat-RNA- and dipeptide repeat protein-driven toxicities, but they provide a direct molecular route by which reduced C9orf72 complex abundance may contribute to previously reported mitochondrial quality-control defects. In disease-relevant settings, proteostasis stress and perturbed phosphorylation signalling could further constrain initiation control and amplify the consequences of reduced SMCR8-FIP200 engagement. More broadly, by defining this molecular interaction and providing mutants that isolate it from other C9orf72 complex functions, our work establishes a mechanistic foundation to rigorously dissect how regulated C9orf72 complex-ULK1/2 complex association shapes mitochondrial quality control in disease-relevant contexts.

## Acknowledgments

We thank members of the Schreiber laboratory for helpful discussions, feedback and suggestions throughout this work. We are grateful to Christopher Green for providing the doxycycline-inducible mKeima cell line and to Huy Le and Melanie Teng for preliminary data. We thank Felix Randow (MRC Laboratory of Molecular Biology) for the FIP200 KO cell line. We also thank the Francis Crick Institute Scientific Technology Platforms (Structural Biology, Cell Services, Screening and Automation, Flow Cytometry, Proteomics, and Bioinformatics and Biostatistics) and the Viral Vector Core Facility for support. We thank James Campbell for advice on statistical analysis; Steven Lim, Sina Namjou, Muhammad Saeed and Ana Agua-Doce for assistance with flow cytometry; Emma Cash for help generating SMCR8-rescued Flp-In cell lines; Rachael Instrell and Ming Jiang for assistance generating the SMCR8 KO cell line; Steven Howell for support with phosphoproteomics; and Molly Strom for assistance with lentivirus production.

The Schreiber laboratory is supported by the Francis Crick Institute, which receives core funding from Cancer Research UK (CC2064), the UK Medical Research Council (CC2064) and the Wellcome Trust (CC2064), and by a UKRI (EPSRC, BBSRC and MRC) Prosperity Partnership award in collaboration with AstraZeneca (EP/X025357/1).

## Author Contributions

A.S. conceived the study with input from J.W. J.W. and C.D. performed all cloning, and expressed and purified proteins. J.W. performed the biochemical analyses and S.K. performed the SPR experiments. J.W. generated cell lines and carried out autophagy flux assays. S.M. and M.S. performed mass spectrometry analyses. J.W. and G.K. analysed the autophagy flux data. J.W. and A.S. assembled the original manuscript draft. J.W., S.K., G.K. and A.S. prepared the figures. J.W. and A.S. reviewed and edited the manuscript with input from all co-authors. A.S. acquired funding.

## Competing Interests

The authors declare no competing interests

## Materials and Methods

### Escherichia coli (E. coli) strains and media

*E. coli* strains (DH5α, BL21-CodonPlus (DE3)-RIL, and DH10Multibac) were cultured in Terrific Broth (TB) medium at 37 °C with shaking at 180 rpm.

### Insect cells and media

Sf9 and High Five insect cells (Thermo Fisher Scientific) were grown in Sf-900 II SFM medium (GIBCO) supplemented with 0.1X Penicillin-Streptomycin-Glutamine (GIBCO) at 27 °C with shaking at 140 rpm.

### Cloning

Genes were either synthesized (Eurofins Genomics) or PCR amplified. Restriction sites and epitope tags were introduced during PCR and PCR products were gel-purified using the GeneJET PCR Purification Kit (Thermo Fisher Scientific) prior to cloning into the pCR-Blunt II-TOPO vector (Thermo Fisher Scientific). Genes encoding proteins for bacterial expression were subcloned into the pET17b-SH-SUMO* plasmid^128^ by restriction enzyme digestion and ligation (**Supplementary Table 1**). Genes encoding proteins or protein complexes for insect cell expression were cloned into pFBDM plasmids^129^ by restriction enzyme digestion and ligation (**Supplementary Table 2**).

For baculovirus-mediated gene delivery to mammalian cells (BacMam), coding sequences were cloned into the BacMam transfer plasmid pBMCL3 by restriction enzyme digestion and ligation. The pBMCL3 plasmid was generated by subcloning the mito-mKeima cassette from pHAGE-mt-mKeima (gift from Richard Youle; Addgene plasmid #131626; RRID:Addgene_131626)^36^ into pBMCL1 (gift from Lei Chen; Addgene plasmid #178203; RRID:Addgene_178203)^130^ after SpeI/PsiI excision and ligation into SpeI/BstZ17I-linearized pBMCL1. The mito-mKeima insert was subsequently replaced with alternative coding sequences to generate the different autophagy reporters (**Supplementary Table 3**).

The tetOperator was removed from pcDNA5/FRT/TO-neo (gift from Jonathon Pines; Addgene plasmid #41000; RRID:Addgene_41000) by SacI digestion followed by ligation to generate pCG_001. The different SMCR8 constructs were then cloned into the multiple cloning site of pCG_001 to generate the plasmids listed in **Supplementary Table 4**.

For lentiviral vector generation, StrepII^2x^-SMCR8-FLAG was cloned into L33.pLT3GEPIR (gift from Johannes Zuber; Addgene plasmid #111177; RRID:Addgene_111177)^131^, yielding pASC_482.

All ligation or In-Fusion reactions were transformed into DH5α competent cells and point mutations were introduced using In-Fusion cloning (TaKaRa Bio). Plasmids were isolated using the GeneJET Plasmid Miniprep Kit (Thermo Fisher Scientific), and all constructs were sequence verified.

### Protein expression in bacteria

Bacterial expression plasmids were transformed into chemically competent BL21-CodonPlus(DE3)-RIL cells (Agilent) and selected on agar plates containing 100 µg/mL ampicillin and 30 µg/mL chloramphenicol. Starter cultures were prepared by inoculating single colonies into 20 mL Terrific Broth (TB) supplemented with antibiotics and incubating overnight at 37 °C with shaking (180 rpm). Cultures were then scaled up to 4 L TB using a starting OD_600_ of 0.01 and grown at 37 °C to an OD_600_ of 0.8-1.0. Cells were cooled on ice for 20 min, and protein expression was induced with 0.5 mM isopropyl β-D-1-thiogalactopyranoside (IPTG), followed by overnight expression at 18 °C with shaking (180 rpm). Cells were harvested by centrifugation at 4,000 rpm (Beckman Coulter J6-MC) for 10 min at 4 °C, and pellets were either processed immediately (see Protein purification below) or flash-frozen in liquid nitrogen and stored at -80 °C.

### Baculovirus generation

MultiBac and BacMam plasmids were transformed into DH10Multibac cells, recovered in 1 mL LB medium (8 h, 37 °C), and plated on low salt agar plates containing ampicillin 100 µg/mL ampicillin, 7 µg/mL gentamycin, 10 µg/mL tetracycline, and 50 µg/mL kanamycin, 0.5 mM IPTG and 300 µg/mL Bluo-Gal for blue-white screening. White colonies were grown overnight in TB medium with antibiotics, and bacmids were extracted via isopropanol precipitation^129^. For baculovirus production, bacmids were transfected into Sf9 cells using GeneJuice Transfection Reagent (Sigma), followed by two rounds of amplification in Sf9 cells to the P3 stage using a multiplicity of infection (MOI) of ∼0.1, following standard protocols.

For baculovirus-mediated gene delivery to mammalian cells (BacMam), pBMCL3-based BacMam transfer vectors encoding the indicated mKeima-based autophagy reporters were used to generate BacMam viruses for transduction of HEK293 cells. The vectors also encoded the vesicular stomatitis virus G protein (VSV-G) to facilitate entry into mammalian cells. Where indicated, Parkin was co-expressed with the mito-mKeima reporter (see “mKeima-based flow cytometry” below).

### Protein expression in insect cells

High Five cells were grown to a density of ∼2 × 10^6^ cells/mL and infected with a P3 baculovirus using a MOI greater than 2. Protein expression was carried out at 27°C for 48-72h and cells were harvested at 4000 rpm (Beckman Coulter J6-MC Centrifuge) for 10 min at 4°C. Pellets were either frozen in liquid nitrogen and stored at -80°C or processed immediately (see Protein purification below).

### Protein purification

#### Cell Lysis

Cell pellets were resuspended in pre-cooled Lysis buffer containing 50 mM Tris-HCl pH 8.5, 300 mM NaCl (for monomeric proteins) or 200 mM NaCl (for protein complexes), 5% glycerol, 2 mM DTT, 4 mM EDTA, 0.2 mM PMSF, 10 µM leupeptin, 10 µM pepstatin A, 1 mM benzamidine, cOmplete EDTA-free protease inhibitor cocktail (Roche; 1 tablet per 50 mL), and Pierce Universal Nuclease (Thermo Fisher Scientific; 6 µL per 100 mL). For GST-tagged proteins, the lysis buffer was adjusted to pH 8.0. For bacterial protein purifications, the Lysis buffer was additionally supplemented with lysozyme (100 µg/mL). Resuspended cell pellets were sonicated and clarified by centrifugation at 20,000 rpm (Beckman Coulter JA-20 rotor) at 4 °C for 1 h.

### Affinity purification

#### StrepTactin purification

For StrepII^2x^-tagged proteins, clarified lysates were loaded onto a StrepTactin Superflow Plus Cartridge (Qiagen) pre-equilibrated in Wash buffer (50 mM Tris-HCl pH 8.0, 200-300 mM NaCl, 5% glycerol, 2 mM DTT; 200 mM NaCl was used for protein complexes and 300 mM for all other proteins). The resin was washed with 20 column volumes (CV) of Wash buffer, and bound proteins were eluted with 5 CV of Wash buffer supplemented with 2.5 mM desthiobiotin (Sigma).

### GST purification

For GST-tagged proteins, clarified lysates were applied to a GSTrap FF column (Cytiva), washed with Wash buffer adjusted to pH 7.6, and eluted with 5 CV of Wash buffer supplemented with 10 mM reduced glutathione (pH 7.6). For GST-ULK1-StrepII^2x^ and GST-ULK2-StrepII^2x^, both Lysis buffer and Wash buffer were additionally supplemented with 50 mM L-arginine and 50 mM L-glutamic acid to enhance protein solubility.

### Tag removal

For constructs requiring tag removal by PreScission (3C) protease (see **Supplementary Tables 1** and **2**), the NaCl concentration was reduced from 300 mM to 200 mM during the final 5 CV of the wash step and maintained at 200 mM in the elution buffer to improve cleavage efficiency. Eluates were incubated overnight with 3C protease at 4 °C at a 1:50 protease-to-protein molar ratio. Following tag removal, samples were either further purified by ion-exchange chromatography or concentrated using Amicon Ultra centrifugal devices (Sigma) prior to size-exclusion chromatography (SEC).

### Anion exchange chromatography

Affinity-purified proteins were diluted with Buffer A (20 mM HEPES-NaOH, pH 8.0, 5% glycerol, 2 mM DTT) to reach a final salt concentration of ∼80-150 mM NaCl (depending on the theoretical pI). Samples were loaded onto a Resource Q column (GE Healthcare) equilibrated in Buffer A (20 mM HEPES-NaOH pH 8.0, 5% glycerol, 2 mM DTT) and the column was washed for 5 CV at the starting salt condition (equivalent to 5% Buffer B, corresponding to ∼35 mM NaCl). Proteins were eluted with a linear NaCl gradient to 700 mM NaCl (equivalent to 100% Buffer B; 20 mM HEPES-NaOH pH 8.0, 700 mM NaCl, 5% glycerol, 2 mM DTT).

### Size exclusion chromatography (SEC)

Proteins purified by affinity chromatography or anion-exchange chromatography were further purified by size-exclusion chromatography (SEC). Samples were loaded onto the appropriate SEC column (see **Supplementary Tables 1** and **2**) pre-equilibrated in SEC buffer (20 mM HEPES-NaOH pH 7.4, 180 mM NaCl, 5% glycerol, 2 mM DTT) and eluted in 1 CV of SEC buffer. When 3C cleavage had been performed, a StrepTactin “passback” column was connected inline downstream of the SEC column to capture the cleaved Strep-tagged moiety and any residual uncleaved protein. Protein-containing fractions were analysed by SDS-PAGE, pooled, concentrated, and snap-frozen.

### Purification of dephosphorylated recombinant C9orf72 complex

The recombinant C9orf72 complex (StrepII^2x^-C9orf72/HA-SMCR8/WDR41-FLAG) was purified as summarised in **Supplementary Table 2**. Because the insect cell–expressed complex was partially phosphorylated, we also prepared a dephosphorylated batch of the complex for better demonstration of its phosphorylation-regulated interaction with the FIP200 Claw domain. The StrepTactin-eluted C9orf72 complex was incubated with 5 mM MgCl_2_ and lambda protein phosphatase (λ-PP; NEB) at 30 °C for 2 h at an approximate 1:20 λ-PP-to-C9orf72 complex molar ratio. Protease inhibitors (1X) were added to minimise protein degradation at this stage. Following dephosphorylation, λ-PP was removed by anion-exchange chromatography, and C9orf72-containing fractions were further purified by SEC on a Superose 6 column. Peak fractions corresponding to the C9orf72 complex were pooled, concentrated and flash-frozen for downstream applications.

### Purification of the C9orf72-SMCR8 subcomplex and the C9orf72 subunit

The recombinant StrepII^2x^-C9orf72-HA-SMCR8 subcomplex was purified as summarised in **Supplementary Table 2**. StrepTactin affinity purification yielded both the StrepII^2x^–C9orf72-HA-SMCR8 complex and StrepII^2x^-C9orf72 alone. The latter was separated from the C9orf72-SMCR8 subcomplex by anion-exchange chromatography. Fractions corresponding to the C9orf72-SMCR8 subcomplex were further purified by size-exclusion chromatography on a Sephacryl S-300 column. Protein-containing peak fractions were pooled, concentrated and flash-frozen for downstream experiments.

#### *In vitro* pulldown assays

Depending on the affinity tag on the bait protein, pulldowns were performed using anti-FLAG M2 affinity resin (Sigma), anti-HA agarose (Sigma), anti-c-Myc agarose (Thermo Fisher Scientific), or StrepTactin Superflow Plus resin (QIAGEN). Prior to sample addition, resins were sequentially washed with 10 bed volumes (BV) each of 1X TBS (50 mM Tris-HCl pH 7.6, 150 mM NaCl), 0.1 M glycine-HCl pH 3.4, and 1× TBS (two washes), and then equilibrated in Pulldown buffer (20 mM HEPES-NaOH pH 7.5, 150 mM NaCl, 2 mM DTT). Bait and prey proteins were mixed at a 1:2 molar ratio and incubated with resin for 15 min at room temperature. Beads were washed four times with Pulldown buffer and bound proteins were eluted in Pulldown buffer containing either 2.5 mM desthiobiotin, 300 µg/mL 3X FLAG peptide (Sigma), 300 µg/mL c-Myc peptide (GenScript), or 100 µg/mL HA peptide (APExBIO). Eluates were analysed by immunoblotting or SDS-PAGE and visualised using Quick Coomassie stain (Neo Biotech) or SYPRO Ruby protein gel stain (Thermo Fisher Scientific).

### Protein phosphorylation prior to pulldown

For phosphorylation reactions, target proteins were incubated with the indicated kinase(s) at a 50:1 substrate-to-kinase molar ratio in Pulldown buffer supplemented with 5 mM MgCl₂, 1 mM ATP, and 1X PhosSTOP phosphatase inhibitor cocktail (Roche) at 37 °C for 45 min. When phosphorylation was required for only a subset of pulldown components, apyrase (NEB) was added after completion of the phosphorylation reaction and samples were incubated at 30 °C for 1 h to deplete residual ATP prior to addition of the remaining protein(s).

### Protein dephosphorylation experiments

For dephosphorylation, proteins were incubated with λ-PP in Pulldown buffer at 30 °C for 1 h at an approximate 1:20 λ-PP-to-substrate molar ratio.

### Surface plasmon resonance (SPR) experiments

SPR experiments were performed on a Biacore S200 instrument (Cytiva). A Strep-Tactin XT-coated CM5 sensor chip was prepared using the Twin-Strep-tag Capture Kit (IBA Lifesciences) according to the manufacturer’s instructions. Measurements were carried out at 25 °C in 1X HBS-P^+^ running buffer (10 mM HEPES-NaOH, pH 7.4, 150 mM NaCl, 0.05% Tween 20; Cytiva) supplemented with 0.5 mM TCEP. The untagged extended C-terminal region of FIP200 (residues 1278-1594) was buffer-exchanged into the same running buffer prior to the SPR experiments.

For binding measurements, different C9orf72 complexes (WT and mutants listed in **Fig. 4f**) were captured via the N-terminal StrepII^2x^-tag on the C9orf72 subunit at a concentration of 25 nM for 90 s at a flow rate of 30 μl/min, yielding surface densities of ∼200-800 response units (RU). A dilution series of FIP200 was then injected over the C9orf72 complex-coated surfaces at 30 µl/min with a contact time of 120-180 s and a dissociation time of 420 s. C9orf72 complexes were subsequently removed from the surfaces by two consecutive 60 s injections of regeneration solution (3 M guanidine hydrochloride) supplied with the Twin-Strep-tag Capture Kit (IBA Lifesciences).

Data were processed using Biacore S200 Evaluation Software (Cytiva). Raw sensorgrams were subjected to double-reference subtraction and baseline alignment. Steady-state response levels were determined from the processed sensorgrams by averaging the signal over 5 s at the end of each injection. Responses were replotted against the FIP200 dimer concentration in Prism 10 (GraphPad). Equilibrium dissociation constants (*K_d_*) were determined by nonlinear least-squares fitting in Prism using a hyperbolic binding equation. Reported *K_d_* values are based on n = 3 to n = 7 independent experiments, depending on the C9orf72 complex. Examples of processed sensorgrams shown in **Extended Data Fig. 4c-k** were also replotted in Prism10.

### Hydrogen-deuterium exchange (HDX) mass spectrometry

The FIP200^CTR^-CSW^SMCR8-LL:S>D/T>E^ complex and CSW^SMCR8-LL:S>D/T>E^ alone were diluted to a final concentration of 10 µM. Deuterium labelling was initiated by diluting 5 µL of protein in 40 µl of D_2_O buffer and allowing exchange to proceed at RT for 3, 30, 300 and 3000 seconds in triplicate. The labelling reaction was quenched by addition of ice-cold 2.4% v/v formic acid in 2 M guanidinium hydrochloride, immediately followed by snap-freezing in liquid nitrogen. Quenched samples were stored at -80°C prior to analysis.

The quenched protein samples were rapidly thawed and proteolytically digested using an Enzymate BEH immobilized pepsin column, 2.1 x 30 mm, 5 µm (Waters) at 200 µL/min for 2 min. Generated peptides were captured on a 2.1 x 5 mm C18 trap column (Acquity BEH C18 Van-guard pre-column, 1.7 µm, Waters) and subsequently eluted over 12 min using a 5-36% gradient of acetonitrile in 0.1% v/v formic acid at 40 µL/min. Peptide separation was achieved using reverse-phase chromatography (Acquity UPLC BEH C18 column 1.7 µm, 100 mm x 1 mm, Waters). Mass spectra were acquired on a Cyclic mass spectrometer (Waters) operated in positive ion mode over a m/z of 300 to 2000, using standard electrospray ionization (ESI) source (source temperature 80°C; spray voltage 3.0 kV). Lock mass calibration was applied using [Glu1]-fibrino peptide B (50 fmol/µL).

Peptide identification was performed by MSe^132^ using an identical gradient of increasing acetonitrile in 0.1% v/v formic acid over 12 min. The resulting MSe data were analyzed using Protein Lynx Global Server software (Waters) with an MS tolerance of 5 ppm. Mass analysis of the peptide centroids was performed using DynamX sotware (Waters). Only peptides with confidence scores greater than 6.4 were considered. Initial peptide assignments were generated automatically by the DynamX software, and all peptides (deuterated and non-deuterated) were subsequently manually verified for all time points to confirm correct charge states, retention times, and the presence of overlapping peptides.

Deuterium incorporation was not corrected for back-exchange and represents relative, rather than absolute deuterium uptake. Changes in H/D amide exchange for individual peptides may reflect exchange at one or multiple backbone amide positions. All labelling reactions were prepared simultaneously, and all samples were analysed on the mass spectrometer on the same day.

### Mammalian cell culture

The Flp-In T-REx 293 cell line was kindly provided by Simon Boulton (Francis Crick Institute). Wild-type HEK293T cells and the corresponding *FIP200*-KO derivative were obtained from Felix Randow (MRC Laboratory of Molecular Biology). The parental HEK293T D8Cas9 cell line was obtained from the Cell Services Science Technology Platform (STP) at the Francis Crick Institute and engineered to express Cas9 nuclease under a doxycycline-inducible promoter.

All cell lines were routinely cultured in DMEM (Gibco) supplemented with 10% fetal bovine serum (Gibco) and 1% penicillin–streptomycin (Sigma) at 37 °C in 5% CO₂. Cells were discarded after 15 passages.

### Generation of *SMCR8*-KO Flp-In T-REx 293 cells

To generate the sgRNA pool the three sgRNA guides (**Supplementary Table 5**) targeting the first exon of SMCR8 were resuspended in TE buffer and combined at a 1:1:1 ratio to yield a 3 µM sgRNA solution. SpCas9-2NLS (Synthego) was resuspended in Opti-MEM medium to yield a final concentration of 3 µM. RNP complexes were assembled by mixing 1.3 µL sgRNA pool, 1 µL SpCas9-2NLS, 1 µL Cas9 Plus Reagent (Invitrogen), and 25 µL Opti-MEM, followed by 5-10 min incubation at room temperature. A transfection solution was prepared separately (25 µL Opti-MEM + 1.5 µL Lipofectamine CRISPRMAX Transfection Reagent, 5 min incubation) and mixed with the RNP complexes in a 1:1 ratio. 1 x 10^5^ Flp-In T-REx 293 cells at ∼70% confluency were trypsinised, resuspended in 500 µL DMEM, mixed with the combined RNP-transfection solution, and plated in one well of a 24-well plate. After 24 h, the medium was replaced with fresh DMEM. Upon reaching confluency, cells were single-cell sorted into 96-well plates, and resulting clones were validated by Western blotting and amplicon sequencing using the Francis Crick Advanced Sequencing Facility.

### Generation of *SMCR8*-KO HEK293T D8Cas9 cell line

A crRNAs pool was prepared by combining five crRNAs (**Supplementary Table 6**) targeting the first exon of the SMCR8 gene. The crRNAs were mixed with tracrRNA (Dharmacon) at a 1:1 molar ratio and diluted to a final guide RNA concentration of 200 nM in phenol red–free HBSS (Gibco). The resultant guide RNA complexes were delivered using Lipofectamine 2000 (Invitrogen) prepared in Opti-MEM (Gibco) at a 1:50 ratio and incubated with cells according to the manufacturer’s instructions. Cells were seeded at a density of 5 × 10⁴ cells/mL in DMEM supplemented with 1 µg/mL doxycycline (Takara) to induce Cas9 expression. Following transfection with 20 nM guide RNA-transfection mix, cells were cultured at 37 °C for four days, after which all wells were pooled and expanded. Single-cell clones were isolated by single-cell cell sorting and screened for *SMCR8*-KO.

### Stable cell line generation

To reintroduce wild-type SMCR8 and SMCR8 mutants into *SMCR8*-KO Flp-In T-REx 293 cells, 3-5 × 10⁶ cells were seeded in 10 cm dishes. At ∼60% confluency, cells were transfected using Lipofectamine 3000 (Thermo Fisher Scientific) with 4.5 µg pOG44 and 0.5 µg of the different pcDNA5/FRT plasmids listed in **Supplementary Table 4**. After 24 h, the medium was replaced. At 48 h post-transfection, cells were re-seeded into fresh 10 cm dishes at 3.5 × 10⁶ cells per dish. Stable integrants were selected in medium supplemented with 15 µg/mL blasticidin (Merck) and 600 µg/mL G418 (Gibco). Selection medium was replaced every 2-3 days, and colonies were monitored for 2-3 weeks until foci became visible.

For the *SMCR8*-KO HEK293T D8Cas9 cells StrepII^2x^-SMCR8-FLAG was introduced by lentiviral transduction. Third-generation lentiviral particles encoding StrepII^2x^-SMCR8-FLAG were produced in HEK293T cells by co-transfecting the transfer plasmid pASC_482 together with the packaging plasmids pLP1 (Invitrogen) and pLP2 (Invitrogen) and the envelope plasmid pVSV-G (System Biosciences). HEK293T cells were cultured in DMEM supplemented with 10% FBS (Biosera) and 1% penicillin–streptomycin (Gibco) and transfected with the packaging plasmids using Lipofectamine 2000 (Invitrogen) in Opti-MEM I (Gibco). Sixteen hours after transfection, the medium was replaced with fresh complete DMEM. Viral supernatants were harvested at 24 and 48 hours post-transfection, cleared by filtration through a 0.45 µm filter (Millex-HV Millipore). Viruses were then concentrated by polyethylene glycol (PEG) precipitation: PEG8000 and NaCl were added to the clarified supernatant to final concentrations of 8% and 80 mM, respectively, followed by incubation at 4 °C for at least 4 hours. Viral particles were pelleted by centrifugation at 3,000 g for 40 minutes, resuspended in PBS to achieve an approximately 50-100-fold concentration, and either used immediately or stored at -80 °C until further use.

Cells were seeded in six-well plates at 4 × 10⁵ cells per well and transduced at a MOI of 1. 48 hours post-transduction, selection was initiated with 0.7 µg/mL puromycin (Sigma), with medium refreshed every two days until resistant cells reached confluency. Cells were subsequently expanded and single-cell clones were isolated by flow cytometry. Expression of SMCR8 was assessed by Western blotting following induction with 1 µg/mL doxycycline.

### Immunoprecipitation experiments

Cells harvested from a T-75 flask were washed once with ice-cold PBS and resuspended in 500-800 µL ice-cold lysis buffer (50 mM Tris-HCl pH 8.0, 180 mM NaCl, 4 mM EDTA, 5% glycerol) supplemented with Pierce Universal Nuclease, 1X protease inhibitor (Thermo Fisher Scientific), and 1X PhosSTOP protein phosphatase inhibitors (Roche). Lysates were sonicated on ice and clarified by centrifugation (15,000 g, 1 h, 4 °C). The clarified supernatant was then incubated with anti-FLAG M2 affinity gel (Sigma) pre-equilibrated in Wash buffer (20 mM HEPES pH 7.8, 180 mM NaCl, 5% glycerol; PhosSTOP was included for phosphoproteomics experiments) for 1-2 h at 4 °C. The resin was washed five times with 10 bed volumes of Wash buffer. Bound proteins were eluted in Wash buffer containing 100 µg/mL 3×FLAG peptide, and eluates were collected for downstream analysis.

### SMCR8 phosphorylation-site mapping by mass spectrometry

HEK293T D8Cas9 cells expressing StrepII^2x^-SMCR8-FLAG were induced with 1 µg/mL doxycycline, and lysates were subjected to anti-FLAG immunoprecipitation using anti-FLAG M2 agarose (see Immunoprecipitation experiments). Eluates were resolved by SDS-PAGE on NuPAGE 4-12% Bis–Tris gels (Invitrogen) in 1X MOPS running buffer (Invitrogen) and stained with SimplyBlue SafeStain (Invitrogen) using the manufacturer’s maximum-sensitivity protocol. The band corresponding to SMCR8 was excised and processed for mass spectrometry analysis.

Briefly, proteins were reduced in 10 mM DTT and then alkylated with 55 mM iodoacetamide. After alkylation, 1 µg of Trypsin (Promega, UK) was added and the proteins digested for overnight at 37 °C in a thermomixer (Eppendorf, Germany), shaking at 800 rpm. The resulting peptides were dried by vacuum centrifugation and phosphopeptide enrichment was carried out using the sequential metal oxide affinity chromatography (SMOAC) strategy with High Select TiO_2_ and Fe-NTA enrichment kits (Thermo Scientific). Eluates were combined, dried by vacuum centrifugation and resuspended in 0.1% trifluoroacetic acid (TFA). Peptides were analysed by liquid chromatography electrospray ionization tandem mass spectrometry (LC-ESI-MS/MS), performed on an Orbitrap Fusion Lumos Tribrid Mass Spectrometer (Thermo Fisher Scientific) equipped with an EasySpray source and UltiMate 3000 RSLCnano UHPLC liquid chromatography system (Thermo Fisher Scientific). The digested protein from each sample was injected in triplicate. 7 μl per injection was loaded onto a 2 mm by 0.3 mm Acclaim PepMap C18 trap column (Thermo Fisher Scientific) in 0.1% TFA at 15 μl/min before the trap being switched to elute at 0.25 μl/min through a 50 cm by 75 μm EASY-Spray C18 column. A ∼90′ gradient of 2 to 20% B over 57′ and then 20 to 40% B over 25′ was used followed by a short gradient to 90% B held for 7’ and back down to 2% B and a 20′ equilibration in 2% B [A = 2% acetonitrile (ACN), 5% DMSO, 0.1% formic acid (FA); B = 75% ACN, 5% DMSO 0.1% FA]. The Orbitrap Fusion Lumos was operated in “Data Dependent Acquisition” mode with the MS1 full scan at a resolution of 120,000 FWHM, followed by as many subsequent MS2 scans on selected precursors as possible within a 3-s maximum cycle time. MS1 was performed in the Orbitrap with an AGC target of 4 × 105, a maximum injection time of 50 ms, and a scan range from 375 to 1500 m/z. MS2 was performed in the ion trap with a rapid scan rate, an AGC target of 2 × 103, and a maximum injection time of 300 ms. Isolation window was set at 1.2 m/z, and 38% normalized collision energy was used for HCD. Dynamic exclusion was used with a time window of 40 s.

All acquired raw mass spectrometric data were processed in MaxQuant (version 1.6.12.10)^133^ for peptide and protein identification; the database search was performed using the Andromeda search engine^134^ against the *Homo sapiens* canonical sequences from UniProtKB. Fixed modifications were set as carbamidomethyl (C) and variable modifications were set as oxidation (M) Acetyl (Protein N-term), and phospho (STY). The estimated false discovery rate was set to 1% at the peptide, protein, and site levels. A maximum of two missed cleavages were allowed. Other parameters were used as preset in the software. The MaxQuant output file PhosphoSTY Sites.txt, an FDR-controlled site-based table compiled by MaxQuant from the relevant information about the identified peptides, was imported into Perseus (v1.4.0.2) for data evaluation.

### Western blot analysis

Adherent cells were washed with ice-cold PBS and lysed in RIPA buffer (Thermo Fisher Scientific) supplemented with 1X PhosSTOP phosphatase inhibitor (Roche), 1X protease inhibitor cocktail, and Pierce Universal Nuclease (Thermo Fisher Scientific). Lysates were incubated on ice for 15-20 min. A small part of the sample was removed to determine protein concentration using the Bradford assay (Thermo Fisher Scientific), and the remainder was mixed with 4X LDS sample buffer supplemented with 100 mM DTT and denatured at 95 °C for 5-10 min. Normalised sample volumes corresponding to 20-30 µg total protein were resolved by SDS-PAGE and transferred to 0.2 µm nitrocellulose membranes (Bio-Rad). Membranes were blocked for 1 h at room temperature in TBS-T (TBS containing 0.05% Tween-20) supplemented with either 5% (w/v) skimmed milk or 5% BSA (for phosphoprotein detection), then incubated overnight at 4 °C with primary antibodies (**Supplementary Table 7**) diluted in TBS-T containing 2% BSA. Membranes were washed three times with TBS-T and incubated for 1 h at room temperature with IRDye-conjugated anti-rabbit or anti-mouse secondary antibodies (LI-COR) diluted 1:10,000 in TBS-T containing 2% BSA. Following three additional washes with TBS-T, membranes were imaged on an Odyssey DLx system (LI-COR).

### Autophagy flux measurements by flow cytometry

Autophagic flux was monitored using monomeric Keima (mKeima)-based reporters delivered by BacMam-mediated transduction. Bulk autophagy was measured using cytosolic mKeima and induced using Torin-1 (100 nM, 24 h; Cell Signaling Technology). Mitophagy was assessed with a mitochondrial matrix-targeted reporter (mito-mKeima) bearing an N-terminal cytochrome c oxidase subunit 8 (COX8) presequence^135^. Parkin-independent mitophagy was induced with ivermectin (16 µM, 24 h; Sigma) and deferiprone (DFP; 1 mM, 48 h; Sigma), whereas Parkin-dependent mitophagy was triggered with oligomycin (Calbiochem)/antimycin A (Sigma) (O/A; 10 µM each; 6 h) or carbonyl cyanide m-chlorophenylhydrazone (CCCP; 5 µM, 6 h; Sigma). For Parkin-dependent conditions, BacMam viruses were used to co-express Parkin together with mito-mKeima. Lysophagy was monitored using an LGALS3–mKeima reporter^111^ and was induced with L-leucyl-L-leucine methyl ester (LLOMe; 1 mM) for 1 hour, followed by washout and an 8-hour recovery in fresh medium. The list of mKeima reporter constructs used for monitoring autophagy is provided in **Supplementary Table 3**.

BacMam transduction was performed in 12-well plates by seeding 1.5-2.0 × 10⁵ cells per well and transducing the following day with 0.5 mL BacMam virus (negative controls received 0.5 mL Sf-900 II SFM). After 8 h, the virus-containing medium was replaced with fresh DMEM. Treatments were initiated 1-2 days after BacMam transduction to allow robust expression of the mKeima reporters while ensuring that cells remained sub-confluent at the time of analysis. Specifically, longer treatments (Torin-1, ivermectin and deferiprone) were started approximately 18 hours after BacMam transduction, whereas shorter treatments (O/A, CCCP and LLOMe) were initiated approximately 40 hours after BacMam transduction. For flow cytometry, cells were trypsinised, pelleted (300 g, 3 min, 4 °C), and resuspended in DMEM containing 5 µg/mL DAPI (Sigma). Cell suspensions were filtered through 5 mL polystyrene round-bottom tubes fitted with cell-strainer caps (Falcon) and analysed on a ZE5 cell analyser (Bio-Rad) using the gating strategy outlined in **Extended Data Fig. 6b**. The following excitation/emission settings were used: DAPI was excited with a 355 nm laser and detected using a 460/22 nm bandpass (BP) emission filter. mNeonGreen was excited with a 488 nm laser and detected using a 525/35 nm BP emission filter. mKeima fluorescence was collected using a 615/24 nm BP emission filter following excitation at 405 nm (neutral form) or 561 nm (acidic form). Data were analysed as described in^105^. Flow cytometry data were analyzed using FlowJo v10.10.0 (BD Biosciences). A ratiometric parameter was generated for each cell using the "Derived Parameters" function. This was calculated as the ratio of the fluorescence emission intensity upon 561 nm excitation to that upon 405 nm excitation. For each mKeima-positive population, the median 561/405 ratio was extracted using the “Add Statistic” function. To quantify treatment-induced autophagic flux for each cell line, the median 561/405 ratio of the treated sample was expressed as fold-change relative to its untreated control.

### Data analysis and statistics

For each biological replicate, the single-cell mKeima excitation ratio (561/405) was calculated as the 615 nm emission intensity upon 561 nm excitation divided by that upon 405 nm excitation, and summarised as the median 561/405 ratio. Within each independent experiment, three biological replicates per cell line × treatment condition were aggregated using the geometric mean of the three medians (equivalent to averaging on the log scale), yielding one value per condition per experiment. These log-transformed values were analysed using a linear regression model with cell line, treatment, and their interaction, while accounting for systematic differences between experimental repeats (experiment included as a blocking factor). Estimated marginal means were obtained and Wald tests were used to evaluate prespecified contrasts, reported as log-fold changes (LFCs). For plots showing fold change (treated/untreated), treated:untreated log-ratios were computed within each experiment and analysed analogously, with Wald tests used to compare treatment effects between cell lines. *P* values were adjusted for multiple comparisons using Holm’s method (family-wise error control). Bars show mean ± s.e.m. across independent experiments; dots represent individual experiments (each derived from biological triplicates). Significance is denoted as *P* < 0.05 (*), *P* < 0.01 (**), *P* < 0.001 (***) and *P* < 0.0001 (****).

Cell lines expressing mNeonGreen-tagged SMCR8 were plotted together with cell lines lacking mNeonGreen expression. However, statistical testing was restricted to specified comparisons indicated in the figures and did not involve indiscriminate comparisons across all cell lines.

### Structure predictions

Structural predictions were generated using the AlphaFold web server^99^. The AlphaFold 3 (AF3) model of the FIP200^CTR^ in complex with an SMCR8-derived peptide comprising both FIR motifs was computed using FIP200 residues 1391-1594 and SMCR8 residues 421-540, with serines 471 and 516 modeled in their phosphorylated states. For clarity, only FIP200 residues 1429-1594 and SMCR8 residues 462-526 are shown in **Fig. 4h**. Model confidence was evaluated using the predicted Inter-residue Distance Difference Test (pIDDT) scores provided by the server.

## Notes

### Competing Interest Statement

The authors have declared no competing interest.

