## Supplemental Data for "Phosphoregulated SMCR8-FIP200 interaction connects the ALS/FTD-linked C9orf72 complex to autophagy initiation and mitochondrial quality control"

### Extended Data Figure 1

**a**

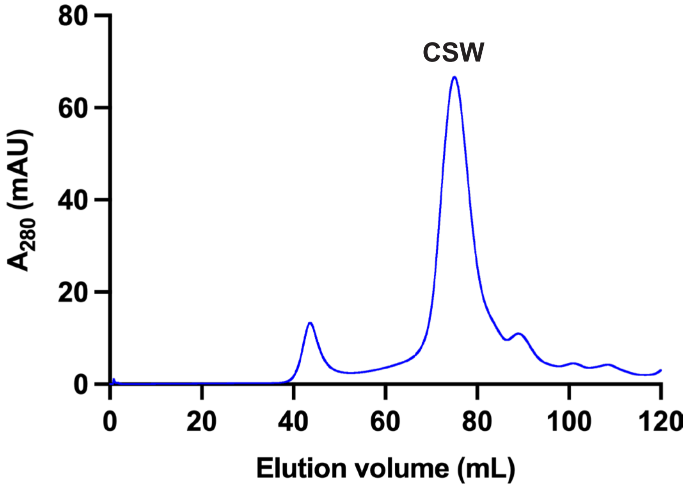

#### Extended Data Figure 2

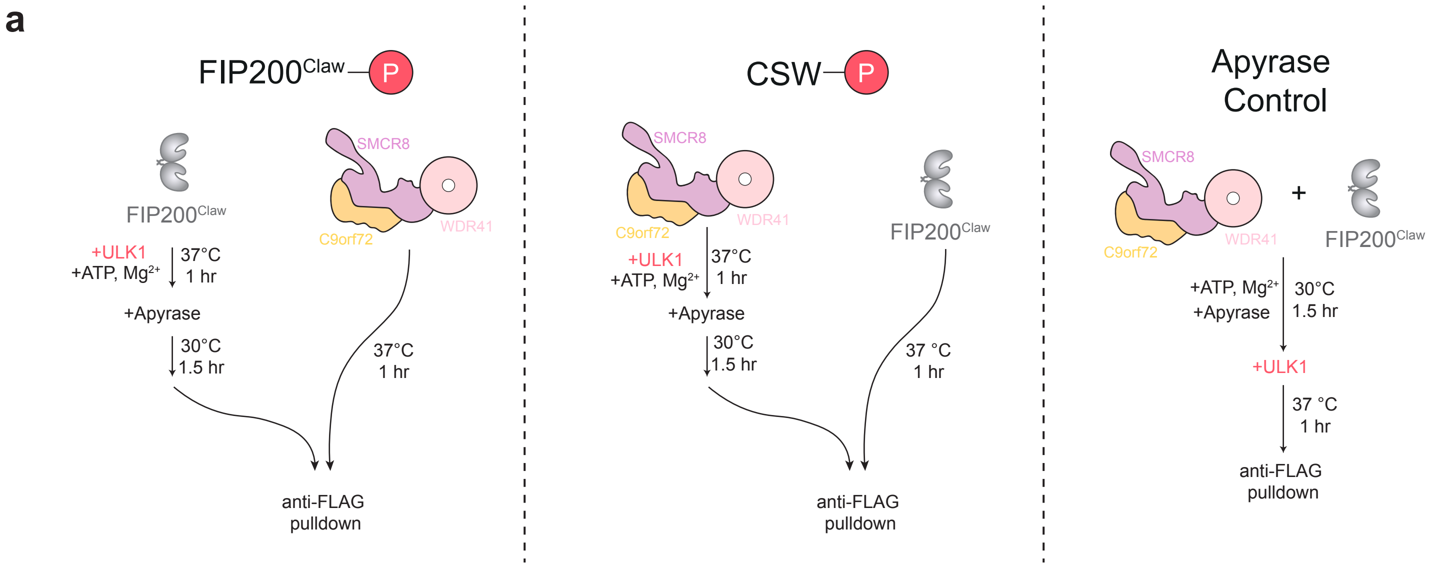

### Extended Data Figure 3

**a**

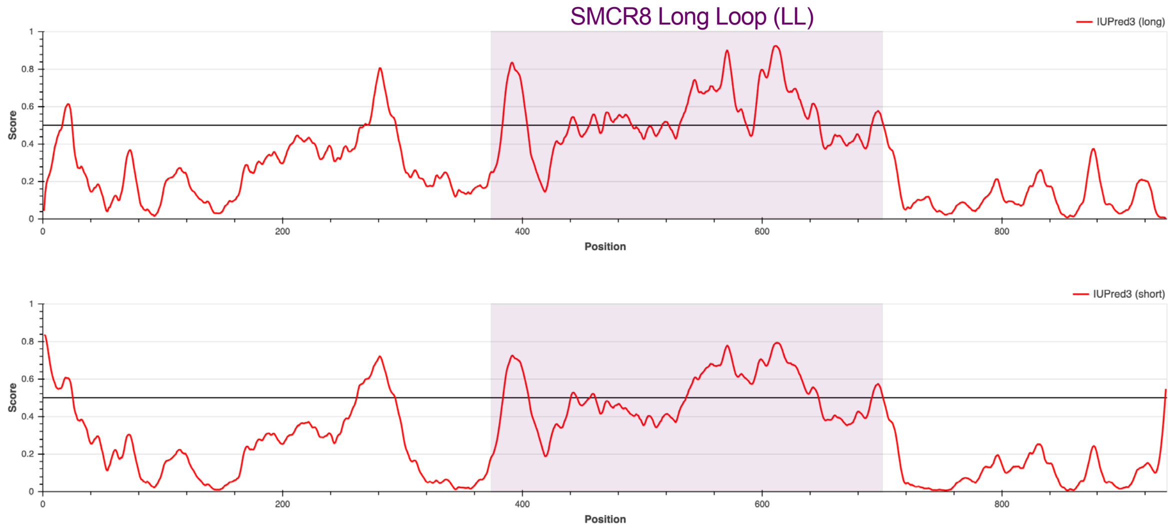

### Extended Data Figure 4

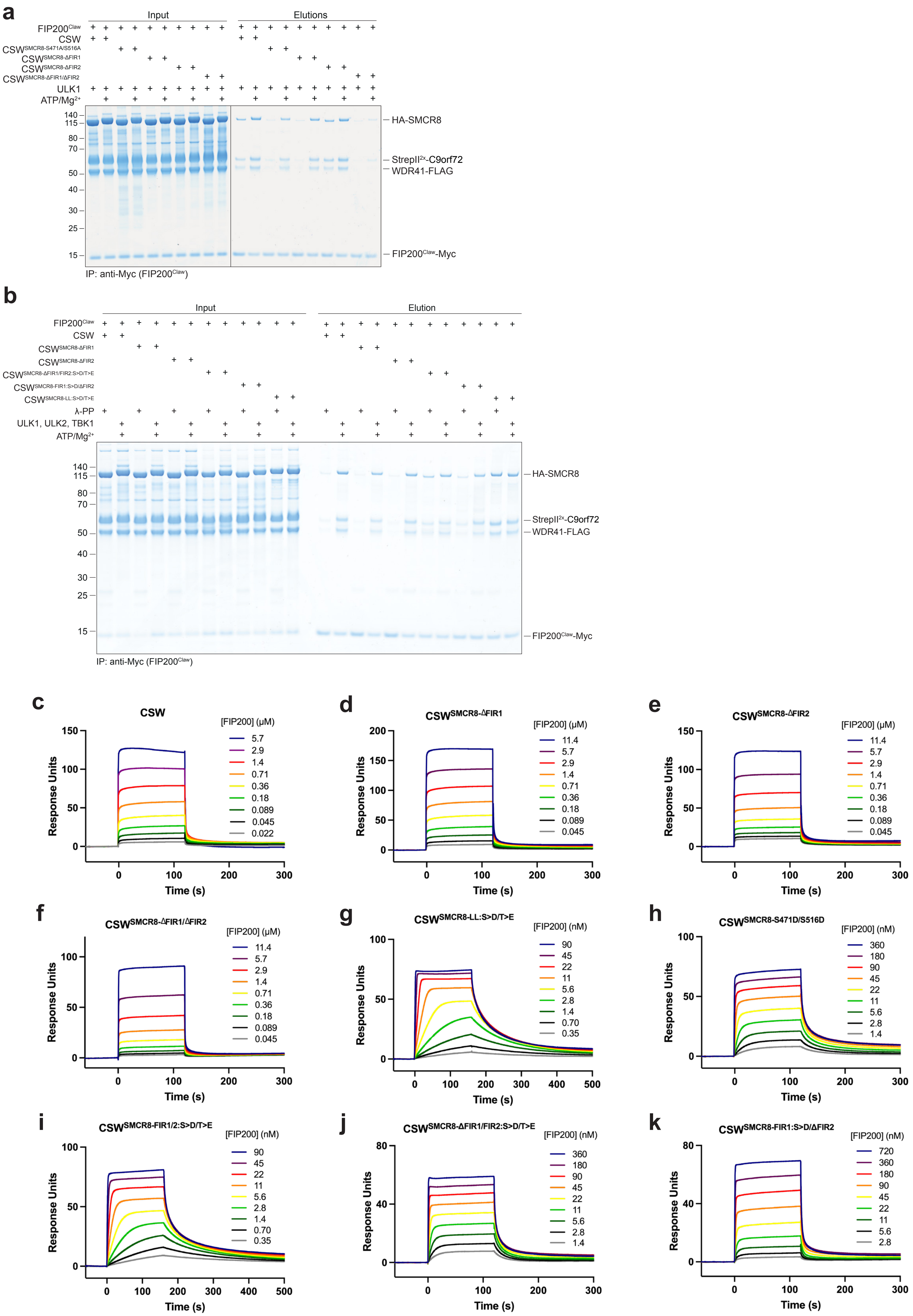

### Extended Data Figure 4

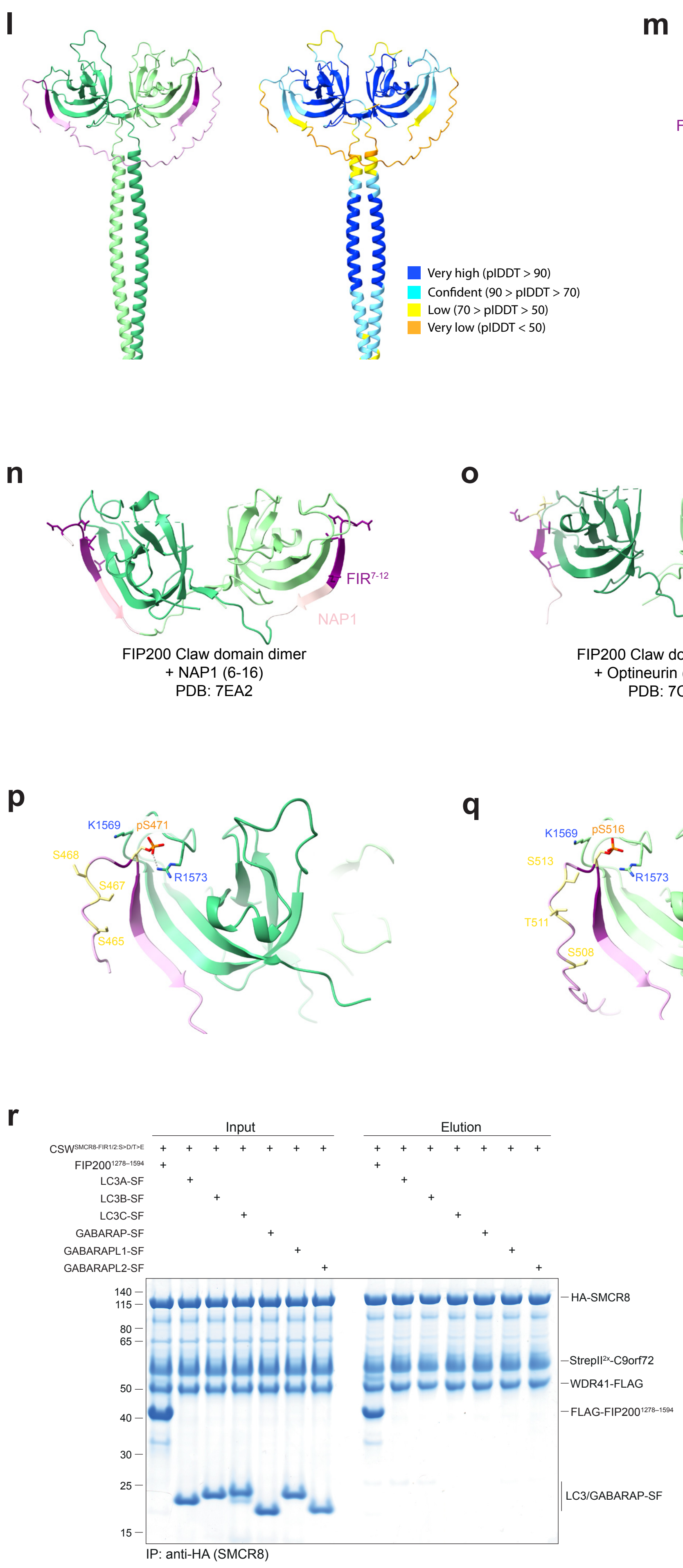

### Extended Data Figure 5

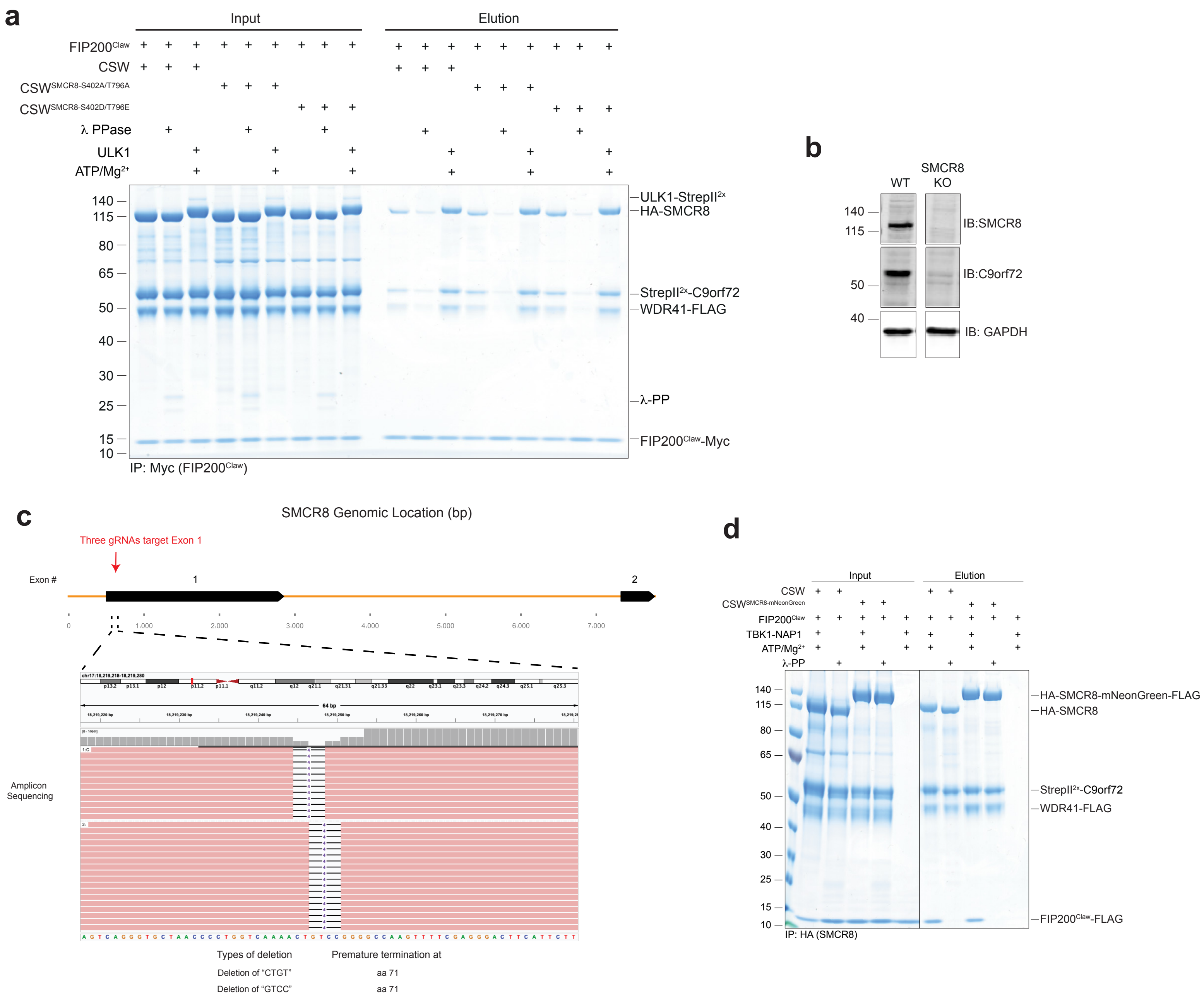

### Extended Data Figure 6

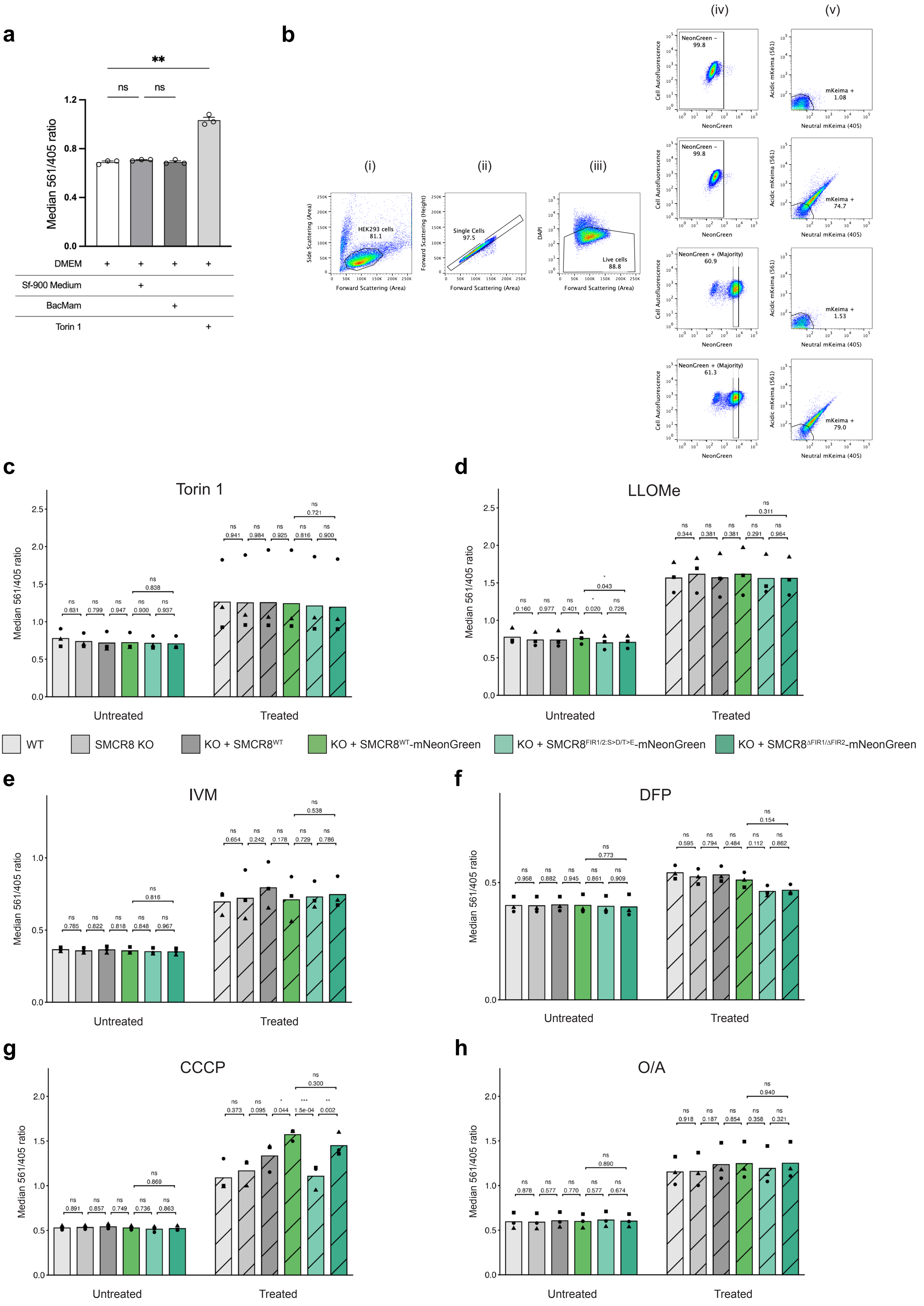

**Extended Data Fig. 1:** Purification of recombinant C9orf72 complex.

**a**, Size-exclusion chromatography (SEC) profile of the recombinant C9orf72 complex. The complex was purified on a HiLoad 16/600 Superose 6 column, and elutions were monitored by UV absorbance at 280 nm. The indicated main peak corresponding to the CSW complex was pooled and used for downstream experiments.

**Extended Data Fig. 2.** ULK1-mediated phosphorylation of the C9orf72 complex strengthens binding to the FIP200 Claw domain.

**a**, Schematic overview of the workflow used in **Fig. 2e**. The C9orf72 complex (CSW) and the FLAG-tagged FIP200 Claw domain ($FIP200^{Claw}$) were individually phosphorylated using substoichiometric amounts of recombinant ULK1. ATP was depleted using apyrase prior to combining the indicated phosphorylated and non-phosphorylated proteins. To verify efficient ATP depletion by apyrase, CSW and $FIP200^{Claw}$ were incubated with $ATP/Mg^{2+}$ and apyrase before ULK1 addition. $FIP200^{Claw}$ was immobilized on anti-FLAG agarose, and after washing, bound proteins were eluted.

**Extended Data Fig. 3.** The intrinsically disordered long loop of SMCR8 mediates FIP200 Claw domain binding.

**a**, Intrinsic disorder prediction of *H. sapiens* SMCR8 using IUPred3. Scores above 0.5 indicate regions predicted to be intrinsically disordered. The analysis was performed using the long- and short-disorder prediction modes of IUPred3 (top and bottom, respectively).

**Extended Data Fig. 4:** Two phosphoregulated FIR motifs in the SMCR8 long loop mediate high-affinity binding to the FIP200 Claw domain.

**a**, Two FIR motifs in the SMCR8 long loop regulate binding to the FIP200 Claw domain. C9orf72 complexes (wild type and indicated mutants) were either phosphorylated by ULK1 (1:20 relative to the C9orf72 complex) or mock-treated by incubation with ULK1 under identical conditions but without $ATP/Mg^{2+}$. C9orf72 complexes were then incubated with Myc-tagged FIP200 Claw domain and captured on Myc resin. After washing, bound proteins were eluted, and input (post-treatment) and elution fractions were analysed by SDS-PAGE followed by Coomassie staining. **b**, Two FIR motifs in the SMCR8 long loop and their phosphorylation regulate binding to the FIP200 Claw domain. Recombinant C9orf72 complexes (wild type and indicated mutants) were either dephosphorylated with lambda phosphatase (λ-PP), or phosphorylated with a mix of recombinant ULK1, ULK2 and TBK1 at a highly substoichiometric ratio (1:20 relative to the C9orf72 complex). The different C9orf72 complexes were then incubated with Myc-tagged FIP200 Claw domain and captured on Myc resin. After washing, bound proteins were eluted, and input (post-treatment) and elution fractions were analysed by SDS-PAGE followed by Coomassie staining. **c-k**, Representative SPR sensorgrams showing binding of the extended C-terminal region of FIP200 (aa 1278–1594) to different C9orf72 complex mutants. C9orf72 complexes were immobilised on a SPR chip via TwinStrep-tag capture and a dilution series of FIP200 was injected over the ligand-coated surface at the concentrations indicated in the figures. Data shown are from a single experiment for each C9orf72 complex variant. **l**, The AF3 model of the FIP200 C-terminus in complex with the FIR motif-containing segment of SMCR8, as shown in **Fig. 4h** (left), is colour-coded by the corresponding pIDDT values (right). Confidence is indicated by pIDDT scores, with values >70 (light blue) or >90 (dark blue) corresponding to high and very high confidence, respectively. **m-o**, Structure-based alignment of the AF3 model shown in **Fig. 4h** and **Extended Data Fig. 4l**. Shown in **(m)** is the FIP200 Claw domain from the AF3 model (residues 1490–1594). In **(n)** this model is aligned with the previously determined structure of the FIP200 Claw domain fused to a FIR motif-containing peptide derived from the TBK1 adaptor NAP1 (residues 6–16; PDB: PDB: 7EA2[61]). In **(o)**, it is aligned with the FIP200 Claw domain in complex with a phosphorylated FIR motif-containing peptide derived from the autophagy cargo adaptor Optineurin (residues 173–183; PDB: 7CZM[60]). The two FIP200 Claw domain protomers are shown in different shades of green; FIR motifs are highlighted in dark purple, and phosphorylated serines in the SMCR8 and Optineurin FIR motifs are indicated in light yellow. **p,q**, Close-up views of the FIR1 **(p)** and FIR2 **(q)** binding sites in the AF3 model shown in **Fig. 4h** and **Extended Data Fig. 4l**. In addition to the core FIR motif serines (pS471 and pS516), upstream Ser/Thr residues (highlighted in yellow) may also be phosphorylated and could partially compensate for the absence of S471/S516 phosphorylation. **r**, The phosphomimicking C9orf72 complex mutant in which the core SMCR8 FIR motif serines and the three upstream Ser/Thr residues were substituted with aspartate or glutamate ($CSW^{SMCR8\text{-}FIR1/2:S>D/T>E}$) was mixed with the indicated LC3- and GABARAP-family proteins and captured on anti-HA agarose. After washing, bound proteins were eluted, and input and elution fractions were analysed by

SDS-PAGE followed by Coomassie staining. A longer FIP200 C-terminal fragment ($FIP200^{1278-1594}$) was included as a positive control.

**Extended Data Fig. 5.** The C9orf72 complex interaction with the ULK1/2 complex depends on the two SMCR8 FIR motifs and their phosphorylation in cells.

**a**, Phosphorylation of S402 and T796 does not regulate FIP200 Claw domain binding. Recombinant C9orf72 complexes (wild type and indicated mutants) were either dephosphorylated with lambda phosphatase (λ-PP) or phosphorylated with recombinant ULK1 at a highly substoichiometric ratio (1:20). The different C9orf72 complexes were then incubated with Myc-tagged FIP200 Claw domain and captured on Myc resin. After washing, bound proteins were eluted, and input (post-treatment) and elution fractions were analysed by SDS-PAGE followed by Coomassie staining. **b**, Validation of the *SMCR8*-KO cell line. *SMCR8* was knocked out in HEK293 Flp-In T-REx cells using CRISPR/Cas9. Cell lysates from wild-type and *SMCR8*-KO cells were analysed by immunoblotting (IB) with the indicated antibodies to confirm loss of SMCR8. Wild-type HEK293 Flp-In T-REx cells served as a positive control. Consistent with previous reports, C9orf72 protein levels were reduced in *SMCR8*-KO cells[16]. **c**, Validation of the *SMCR8*-KO cell line by amplicon sequencing. Amplicon sequencing identified two CRISPR-induced 4-nucleotide deletions, each causing a frameshift and premature termination at amino acid 71. **d**, C-terminal mNeonGreen tagging of SMCR8 does not affect FIP200 Claw domain binding. Purified recombinant C9orf72 complexes containing HA-tagged SMCR8 either lacking or carrying a C-terminal mNeonGreen-FLAG tag were treated with λ-PP or phosphorylated with TBK1 before immobilisation on anti-HA agarose, and then incubated with the FIP200 Claw domain to assess whether the tag affects binding. After washing, bound proteins were eluted, and input and elution fractions were analysed by SDS-PAGE followed by Coomassie staining.

**Extended Data Fig. 6.** The SMCR8-FIP200 interaction differentially regulates selective autophagy pathways.

**a**, BacMam transduction does not measurably alter autophagy. Flp-In T-REx 293 cells expressing a doxycycline-inducible mKeima reporter were incubated with either a control BacMam virus (not encoding a transgene) or the corresponding virus medium (Sf-900 II SFM) to assess potential effects of viral transduction and viral medium on autophagy, respectively. Torin 1 was included as a positive control for autophagy induction. Autophagy was quantified using the mKeima 561/405 excitation ratio. Bars represent the mean ± SEM of triplicates from a representative experiment. Statistical significance was assessed by one-way ANOVA followed by Dunnett's multiple comparisons test to compare treatment groups to the vehicle/medium control. $P < 0.05$ (*), $P < 0.01$ (**), $P < 0.001$ (***), and $P < 0.0001$ (****). **b**, Flow cytometry workflow for autophagy analysis. HEK293 cells were distinguished from debris based on forward- and side-scatter characteristics (i). Singlets were selected using FSC-A vs FSC-H gating (ii), and live cells were selected based on low DAPI uptake (iii). To evaluate effects of changes in the SMCR8-FIP200 interaction, only cells expressing comparable levels of mNeonGreen-tagged SMCR8 (WT or mutants) were analysed (iv). Cells with higher emission upon 405- and 561-nm excitation relative to non-transduced controls, were classified as mKeima-positive and taken forward for ratiometric autophagy flux analysis. The "mKeima+" gate was positioned such that ~99% of untransduced cells fell inside the gate (negative); all cells outside this gate were considered mKeima-positive (v). Approximately 20,000 mKeima-positive cells per sample were acquired for analysis. **c-h**, Raw median 561/405 ratios underlying **Fig. 6e-j**. For each sample, the single-cell mKeima excitation ratio (561/405) was calculated as the 615 nm emission intensity upon 561 nm excitation divided by that upon 405 nm excitation, and the median ratio was determined for each condition. Panels show the raw median 561/405 ratios (a.u.) for untreated and treated samples across the indicated cell lines for the same perturbations as in **Fig. 6e-j** (c corresponds to e; d to f; e to g; f to h; g to i; h to j). Bars show mean ± s.e.m. from three independent experiments (cell-line colour key shown below panels c and d); dots indicate experiments (each performed in biological triplicate per condition).

#### Supplementary Tables

**Supplementary Table 1 |** Overview of recombinant proteins expressed in *E. coli* (BL21-CodonPlus(DE3)-RIL) and purification workflows.

| Plasmid ID | Protein Construct | Purification Strategy |
|---|---|---|
| pJW_310 | SH-SUMO*-OPTN-Myc | StrepTactin > 3C > Superose 6 |
| pASC_701 | SH-SUMO*-FIP200$^{1278\text{-}1594}$-Myc | StrepTactin > 3C > Superdex 75 |
| pJW_135 | SH-SUMO*-FIP200$^{1330\text{-}1594}$-Myc | StrepTactin > 3C > Superdex 75 |
| pJW_136 | SH-SUMO*-FIP200$^{1330\text{-}1493}$-Myc | StrepTactin > 3C > Superdex 75 |
| pJW_201 | SH-SUMO*-FIP200$^{1494\text{-}1594}$ | StrepTactin > 3C > Superdex 75 |
| pJW_200 | SH-SUMO*-FIP200$^{1494\text{-}1594}$-FLAG | StrepTactin > 3C > Superdex 75 |
| pJW_413 | SH-SUMO*-FIP200$^{1494\text{-}1594}$-Myc | StrepTactin > 3C > Superdex 75 |
| pJW_198 | SH-SUMO*-FIP200$^{1494\text{-}1594;\ K1569A,\ R1573E}$-FLAG | StrepTactin > 3C > Superdex 75 |
| pASC_678 | LC3A-SF | StrepTactin > 3C > Superdex 75 |
| pASC_671 | LC3B-SF | StrepTactin > 3C > Superdex 75 |
| pASC_672 | LC3C-SF | StrepTactin > 3C > Superdex 75 |
| pASC_704 | GABARAP-SF | StrepTactin > 3C > Superdex 75 |
| pASC_706 | GABARAPL1-SF | StrepTactin > 3C > Superdex 200 |
| pASC_675 | GABARAPL2-SF | StrepTactin > 3C > Superdex 75 |

**Abbreviations|** SH-SUMO*: $His_6$-2×StrepII$^{2X}$–SUMO* tag; SF: StrepII$^{2X}$-FLAG; 3C: tag cleavage with PreScission protease (HRV 3C).

**Supplementary Table 2 |** Overview of recombinant proteins expressed in High Five insect cells and purification workflows.

| Plasmid ID | Protein Construct | Purification Strategy |
|---|---|---|
| pJW_018 | StrepII$^{2x}$-ATG13/Myc-ATG101 | StrepTactin > ResQ > Superdex 200 |
| pJW_043 | StrepII$^{2x}$-ULK3 | StrepTactin > Superdex 200 |
| pJW_091 | StrepII$^{2x}$-C9orf72/HA-SMCR8 | StrepTactin > ResQ > Sephacryl S-300 |
| pJW_099 | StrepII$^{2x}$-C9orf72/HA-SMCR8/WDR41-FLAG | StrepTactin > Superose 6 |
| pJW_176 | StrepII$^{2x}$-C9orf72/HA-SMCR8$^{\Delta FR}$/WDR41-FLAG | StrepTactin > Superose 6 |
| pJW_177 | StrepII$^{2x}$-C9orf72/HA-SMCR8$^{\Delta LL}$/WDR41-FLAG | StrepTactin > Superose 6 |
| pJW_175 | StrepII$^{2x}$-C9orf72/HA-SMCR8$^{LL:S>D/T>E}$/WDR41-FLAG | StrepTactin > Superose 6 |
| pJW_174 | StrepII$^{2x}$-C9orf72/HA-SMCR8$^{FR:S>D/T>E}$/WDR41-FLAG | StrepTactin > Superose 6 |
| pJW_158 | StrepII$^{2x}$-C9orf72/HA-SMCR8$^{S402D/T796E}$/WDR41-FLAG | StrepTactin > Superose 6 |
| pJW_156 | StrepII$^{2x}$-C9orf72/HA-SMCR8$^{S402A/T796A}$/WDR41-FLAG | StrepTactin > Superose 6 |
| pJW_318 | StrepII$^{2x}$-C9orf72/HA-SMCR8$^{S471A/S516A}$/WDR41-FLAG | StrepTactin > Superose 6 |
| pJW_320 | StrepII$^{2x}$-C9orf72/HA-SMCR8$^{S471D/S516D}$/WDR41-FLAG | StrepTactin > Superose 6 |
| pJW_261 | StrepII$^{2x}$-C9orf72/HA-SMCR8$^{FIR1/2:S>D/T>E}$/WDR41-FLAG | StrepTactin > Superose 6 |
| pJW_316 | StrepII$^{2x}$-C9orf72/HA-SMCR8$^{\Delta FIR1/\Delta FIR2}$/WDR41-FLAG | StrepTactin > Superose 6 |
| pJW_220 | StrepII$^{2x}$-C9orf72/HA-SMCR8$^{\Delta FIR1}$/WDR41-FLAG | StrepTactin > Superose 6 |
| pJW_221 | StrepII$^{2x}$-C9orf72/HA-SMCR8$^{\Delta FIR2}$/WDR41-FLAG | StrepTactin > Superose 6 |
| pJW_032 | StrepII$^{2x}$-FIP200-FLAG | StrepTactin > Superose 6 |
| pJW_055 | StrepII$^{2x}$-FIP200$^{1-640}$-FLAG | StrepTactin > Superose 6 |
| pJW_303 | StrepII$^{2x}$-FIP200$^{641-1329}$-Myc | StrepTactin > ResQ > Superose 6 |
| pJW_167 | GST-ULK1$^{WT}$-StrepII$^{2x}$ | StrepTactin > 3C > Pass back (GST) |
| pJW_168 | GST-ULK2$^{WT}$-StrepII$^{2x}$ | StrepTactin > 3C > Pass back (GST) |

| | | |
|---|---|---|
| pJW_084 | StrepII$^{2x}$-TBK1$^{WT}$/NAP1-Myc | StrepTactin > Superose 6 |
| pJW_238 | StrepII$^{2x}$-TAX1BP1 | StrepTactin > Superose 6 |

**Supplementary Table 3 |** pBMCL3 plasmids encoding mKeima-based autophagy reporters used for monitoring autophagy flux in cells.

| Plasmid ID | Autophagy reporter (promoter) |
|---|---|
| pASC_724 | mKeima (CMV) |
| pASC_886 | mito-mKeima (CMV)/Parkin (CMV) |
| pASC_617 | mito-mKeima (CMV) |
| pASC_1011 | mKeima-LGALS3 (CMV) |

**Supplementary Table 4 |** pcDNA5/FRT plasmids encoding SMCR8 constructs used for generation of stable Flp-In T-REx 293 cell lines.

| Plasmid ID | SMCR8 construct |
|---|---|
| pASC_979 | StrepII$^{2x}$-SMCR8$^{WT}$-FLAG |
| pASC_992 | StrepII$^{2x}$-SMCR8$^{WT}$-mNeonGreen-FLAG |
| pASC_1000 | StrepII$^{2x}$-SMCR8$^{FIR1/2:S>D/T>E}$-mNeonGreen-FLAG |
| pASC_1001 | StrepII$^{2x}$-SMCR8$^{\Delta FIR1/\Delta FIR2}$-mNeonGreen-FLAG |

**Supplementary Table 5 |** The sgRNA sequences used for knocking out the SMCR8 gene in Flp-In T-REx 293 cells.

| **sgRNA sequences** |
|---|
| 5'-U*U*C*CGAGUUCUCUGAGCAGG-3' |
| 5'-A*C*C*CCUGGUCAAAACUGUCC-3' |
| 5'-C*U*C*CCUGCGUAUCAUGUCUG-3' |

Asterisks (*) denote Synthego 5′ terminal sgRNA chemical modifications (2′-O-methyl and phosphorothioate) that increase nuclease resistance.

**Supplementary Table 6 |** The crRNA sequences used for knocking out the SMCR8 gene in HEK293T D8Cas9 cells.

| **crRNA sequences** |
|---|
| 5'-ATCCTCGTCAGAGTGTGTGT-3' |
| 5'-GTTGTCTCGGGAAGAAGGGT-3' |
| 5'-GAGGAGGCCCAATGTTTCAC-3' |
| 5'-GCTCGGGTCCAGCTCATAAG-3' |
| 5'-CGTAGTGGCCTTCACCAAAG-3' |

**Supplementary Table 7 |** List of primary antibodies used in this study.

| **Antigen** | **Catalogue No.** | **Supplier** |
|---|---|---|
| SMCR8 | ab202283 | Abcam |
| C9orf72 | GTX632041 | GeneTex |
| FIP200 | #12436 | Cell signaling technology |
| ULK1 | 8054S | Cell signaling technology |
| Phospho-ULK1 (Ser757) | 6888S | Cell signaling technology |
| MTCO2 | ab110258 | Abcam |
| TOM20 | sc-17764 | Santa Cruz Biotechnology |
| NDP52 | #60732 | Cell signaling technology |
| OPTN | 10837-1-AP | Proteintech |
| TAX1BP1 | #5105S | Cell signaling technology |
| SQSTM1/p62 | ab56416 | Abcam |
| Parkin | sc-32282 | Santa Cruz Biotechnology |
| GAPDH | MAB374 | Merck-Millipore |
| Vinculin | V9131 | Sigma Aldrich |
| mKeima | M126 | MBL Life Science |
| FLAG | F1804 | Sigma |
